# Spatial Transcriptomics Reveals Inflammation and Trans-differentiation States of Acute Myeloid Leukemia in Extramedullary and Medullary Tissues

**DOI:** 10.1101/2024.11.11.622999

**Authors:** Enes Dasdemir, Ivo Veletic, Christopher P. Ly, Andres E. Quesada, Christopher D. Pacheco, Fatima Z. Jelloul, Pamela Borges, Sreyashi Basu, Sonali Jindal, Zhiqiang Wang, Alexander Lazar, Khalida M. Wani, Dinler A. Antunes, Patrick K. Reville, Preethi H. Gunaratne, Robert J. Tower, Padmanee Sharma, Hussein A. Abbas

**Affiliations:** Department of Leukemia, Division of Cancer Medicine, The University of Texas MD Anderson Cancer Center, Houston, TX, USA; Department of Biology and Biochemistry, University of Houston, Houston, TX, USA; Department of Hematopathology, Division of Cancer Medicine, The University of Texas MD Anderson Cancer Center, Houston, TX, USA; Immunotherapy Platform, James P. Allison Institute, The University of Texas MD Anderson Cancer Center, Houston, TX, USA; Department of Pathology, Division of Pathology/Lab Medicine, The University of Texas MD Anderson Cancer Center, Houston, TX, USA; Department of Genitourinary Medical Oncology, Division of Cancer Medicine, The University of Texas MD Anderson Cancer Center, Houston, TX, USA; Department of Surgery, University of Texas Southwestern Medical Center, Dallas, TX, USA; Department of Genomic Medicine, Division of Cancer Medicine, The University of Texas MD Anderson Cancer Center, Houston, TX, USA

## Abstract

Acute myeloid leukemia (AML) is a heterogeneous disease of the bone marrow (medullary) but can also involve extramedullary tissues. While single cell dynamics of AML in suspension are previously explored, a comprehensive spatial transcriptomic assessment in AML remain underexplored. Here, we used Visium spatial transcriptomics to resolve medullary and extramedullary AML environments. We reveal spatial co-localization of monocytes and granulocyte-monocyte progenitors with leukemic populations in the bone marrow, sharing molecular signatures with extramedullary sites. Cell-cell communication via the CXCL12- CXCR4 axis correlated with PI3K/AKT/mTOR signaling in high inflammatory niches. Trans- differentiation states were concentrated in AML-infiltrated regions, with committed-like AML populations present in inflammatory niches and away from the trabeculae, while primitive-like AML cells localized near the endosteal niche. We validated these findings in GeoMx-based Digital Spatial profiling (DSP). Our study applied multimodal spatial transcriptomic approaches to characterize the spatial hierarchy and microenvironmental dynamics of AML differentiation states. We also demonstrated the feasibility of applying Visium-based spatial transcriptomics in decalcified bone tissues.

## Introduction

Acute myeloid leukemia (AML) is a clonal disorder characterized by the presence of immature blasts and arrested differentiation.^1^ Even with the advent of new treatments^2,3^, the overall five-year survival rate remains dismally low at around 30%.^4^ Clinical manifestations of AML are largely attributed to the ability of myeloid neoplastic cells to impair hematopoiesis and disrupt the immune system function in the bone marrow (BM).^5,6^ However, the traditionally view of AML as a BM-centric illness, fails to encapsulate its ability to infiltrate extramedullary (EM) tissues, a condition also known as myeloid sarcoma (MS), which occurs in up to 22% of AML patients.^7^ Interestingly, some AML patients may present with isolated MS without BM involvement.^8^ These EM manifestations underscore the disease’s insidious capacity for dissemination beyond the BM and its adaptability to extra-osseous sites. A thorough investigation of AML’s spatial interactions may reveal the distinct behavior of AML cells in various microenvironments and aid in understanding its adaptability to difference niches.

Recent advances in high-throughput single-cell genomics and transcriptomics have improved our ability to molecularly dissect cellular identities, unravel gene regulatory networks, and characterize cellular states and their interactions within the BM and tumor microenvironment.^9–11^ However, due to tissue dissociation, these approaches result in the loss of spatial context, which is crucial for preserving microenvironmental information.^12^ While spatial proteomic analysis has provided valuable perspectives on mechanisms of AML to evade immune surveillance^13,14^ and revealed subcellular compartments in AML cells,^15^ spatial proteomics can be constrained by a predetermined, targeted approach, limiting its breadth to a fixed set of proteins and favoring validation over novel discovery. In contrast, spatial transcriptomics (ST) has emerged as an investigative tool, capturing a vast array of genes without bias, and facilitating the discovery of biological pathways, molecular profiling, and cellular interactions.^16–19^

Within the array of spatial transcriptomic methodologies^20,21^, the Visium pipeline from 10X Genomics offers comprehensive transcriptome profiling, though its resolution is confined to spots of 55 µm, which restricts its application for single-cell transcriptomics. Nonetheless, it supports high-throughput RNA profiling and facilitates the integration with single-cell RNA sequencing (scRNA-seq) data from cells in suspension, whereas the analysis is enhanced through deconvolution techniques.^22–27^ While the application of Visium-based ST in solid tumors is on the rise,^28–32^ to the best of our knowledge at the time of writing this manuscript, its potential in BM diseases and specifically AML, remains less explored. Also, the rigorous decalcification required for processing and sectioning of BM specimens may compromise RNA integrity, coupled with the perception of BM diseases as ‘liquid’ cancers lacking a defined tissue architecture, may have impeded advancements in this area.

In this study, we performed Visium array-based ST using two different chemistry versions in BM and EM leukemias from two AML patients with tissues collected at the time of diagnosis. We investigated the cellular composition across different BM and EM samples, highlighting spatial heterogeneity. By adapting a median absolute deviation (MAD)-based spot-filtering approach for unique structure metrics and using BM suspension scRNA-seq reference from 16 samples (n=9 healthy and n=7 AML) for spatial deconvolution, we provided a spatial map of medullary and extramedullary leukemia. Our findings reveal the connection between leukemic populations via the CXCL12-CXCR4 axis and the spatial patterns of its downstream targets across the BM-EM axis. Additionally, we showed the involvement of inflammatory and endosteal BM niches in the maturation states of AML populations. We then validated our findings using digital spatial profiling (DSP) in an independent group of AML patients to better delineate the spatial relationship of AML hierarchy to bone structures.

## Materials and methods

This study adheres to the principles outlined in the Declaration of Helsinki. All uses of human material were conducted following written informed consent, which was approved by the institutional review board at The University of Texas MD Anderson Cancer Center.

## Sample preparation

BM and EM tissue biopsies were routinely collected before treatment initiation from 2 patients with newly diagnosed AML. Samples were selected based on sample availability and were formalin-fixed, and BM samples were decalcified using 10% formic acid. Following fixation and decalcification, the samples were embedded in paraffin and stored at room temperature before they were cut into 4-µm sections.

## Clinical immunohistochemistry

For immunohistochemistry (IHC) staining, tissue sections were first deparaffinized and rehydrated in xylene followed by graded alcohols. Endogenous peroxidase activity was blocked using 3% hydrogen peroxide. Heat-induced antigen retrieval was performed in a citrate buffer at 95°C. Following a protein block with 2.5% goat serum, primary antibodies specific to relevant markers (e.g., CD11c, MPO, and CD3e) were applied to the sections and incubated for 1 hour. The primary antibodies were detected using a horseradish peroxidase-conjugated secondary antibody and a DAB substrate. Nuclei were counterstained with hematoxylin. Finally, the stained slides were dehydrated and mounted with coverslips.

## GeoMx Digital Spatial Profiling

An 8x8 core tissue microarray was created from 12 formalin-fixed, paraffin-embedded bone core biopsy samples. Nine cores were selected for DSP characterization. Two cores (areas of illumination [AOI] 1-3) were not included in the present study due to a diagnosis of M6 (acute erythroid) leukemia. GeoMx sample preparation was performed as described previously.^33^ For 3 cores, region of interest (ROIs) were selected both adjacent to and at least 200 µm distal from bone structures to characterize the tumor microenvironment proximal and distal to the bone. The tissue microarray was then stained with anti-CD68 (KP1) antibody, anti-CD34 (QBend/10) antibody, and SYTO13 nuclear stain to identify AML-enriched areas and to segment the ROIs into CD68, CD34, and non-myeloid areas of illumination (AOIs). Oligonucleotide barcodes attached to in situ hybridization probes from the GeoMx whole transcriptome atlas WTA were used to capture RNA expression for over 18,000 genes. The barcodes were cleaved using a UV laser to spatially capture expression in each AOI, then quantified.

RNA data underwent quality assurance using the GeomxTools R package.^34^ In brief probes under the limit of quantification (defined as 2 standard deviations above the geometric mean of the negative control probes for each segment) in were removed from analysis. Probe counts were then normalized using quartile 3 (Q3) normalization and log transformed as recommended by NanoString.^33^ RNA samples from myeloid AOIs were deconvolved *in silico* with CIBERSORTx^35^ using our generated cell types as the reference. To simplify analysis, common myeloid progenitor/lymphoid-primed multipotent progenitor-like and HSC-like categories were combined into primitive-like, basophil-like, dendritic cell-like, and monocyte-like were combined into committed-like, and lymphoid-like and common lymphoid progenitor-like were combined into lymphoid-like. Differences in cell type abundance by proximity to bone were compared using a paired two-sample t-test. All analysis was performed using the R statistical language version 4.3.0.

## Visium ST experimental methods

Formalin-fixed paraffin-embedded (FFPE) tissue from 2 BM and 2 EM leukemia samples were used for spatial transcriptomics analysis. Samples were paraffin-embedded and serially sectioned (thickness 4 μm). FFPE samples were tested for RNA quality with a DV_200_ measurement by 2100 Electrophoresis Bioanalyzer (Agilent; G2939BA). The samples were then processed according to the standard Visium-Cytassist Spatial Gene Expression protocol (CytAssist-enabled) for the v2 assay using Visium CytAssist Reagent Kit, and standard Visium Spatial Gene Expression protocol (Direct placement) for the v1 assay (10X Genomics). The samples to be used for the v1 assay were directly placed on the Visium Spatial Gene Expression slide, which contains spatially unique capture oligonucleotides with 6.5 x 6.5 mm capture area. The samples for the v2 assays were placed on a standard glass slide and processed through Visium CytAssist and transferred to Visium’ 11 x 11 mm capture area slide.

Libraries were cleaned up using SPRI select reagent and quantified using the High Sensitivity DNA Kit run on the Agilent 2100 Bioanalyzer, as well as the KAPA Library Quantification Kit for the Illumina platform (Roche, 7959362001) run on LightCycler 480. The library pool was quantified on the Bioanalyzer and with quantitative polymerase chain reaction and was sequenced using Illumina NovaSeq 6000.

## Visium ST analysis workflow

### Raw read processing with SpaceRanger

Processing of spatial data was conducted using the SpaceRanger (version 2.0) software suite provided by 10X Genomics. For datasets derived from the Visium v1, raw base call (BCL) files were generated. These BCL files were transformed into FASTQ format utilizing the mkfastq function of the SpaceRanger. Datasets obtained from the Visium v2 were directly procured in FASTQ format. These FASTQ files were then aligned to the human reference genome GRCh38, sourced from 10X Genomics, utilizing the count function to map barcoded spots to individual slides. The analytical evaluation involved examining metric summaries and web summary files for each sample to facilitate comparative analysis. To evaluate the performance of each sample, median values of the number of oligonucleotides and genes metrics were calculated.

### Pathology Annotation

H&E-stained slides were digitized (40x) using the Aperio from Leica Biosystems. Clinical IHC-stained slides were scanned at 40x using an Akoya Biosciences Vectra Polaris automated quantitative pathology imaging system. Images were annotated by expert pathologists using the Aperio ImageScope pathology slide viewing software (version 12.4.6). Spots that fall in these annotated regions were identified using the Loupe Browser. The spot barcodes and annotations were exported as a CSV file and added to the metadata of our Seurat objects. These annotations were then visualized using the *SpatialDimPlot* function.

### Spatial MAD

Read count matrices and spatial informations were loaded using the *Load10X_Spatial* function from the *Seurat* R package (version 5.0.3). Mitochondrial content was assessed using the *PercentageFeatureSet* function. The log10 value of genes per unique molecular identifier was calculated as log10(number of genes) / log10(number of oligonucleotides). Initially the median value of mitochondrial read percentage was determined, and maximum and minimum thresholds were set at +3 and –3 times median absolute deviation (MAD), respectively. The same procedure was applied to the log10 values of number of genes detected. Additionally, the +3 MAD was used to define only the maximum threshold for log10(number of oligonucleotides) values. This automated filtering process was applied to each sample individually.

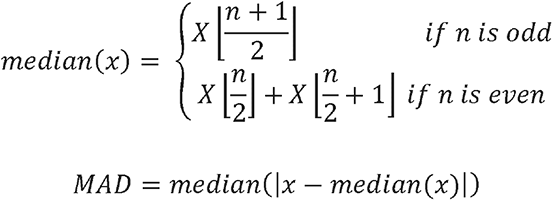

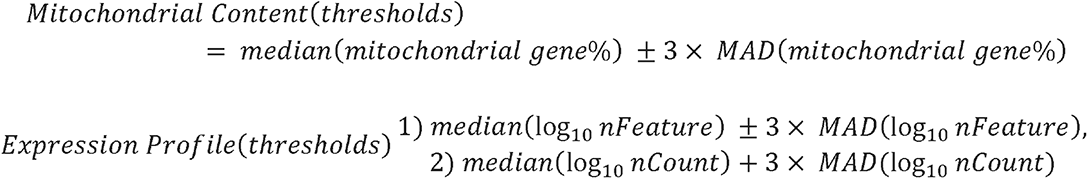

*χ* = Feature of the ST dataset

*n* = number of the values in the feature dataset

*nFeature* = Number of gene captured

*nCount* = Number of oligonucleotide captured

The data was visualized with ggplot2 (version 3.5.0) and ggExtra (version 0.10.1) to confirm the filtering criteria.

### Visium ST data processing

Filtered spots were normalized using the *SCTransform*^36^ function in *Seurat* (version 5.0.3).^37^ The top 5,000 variable genes were then identified using the variance stabilizing transformation method through the *FindVariableFeatures* function. Subsequently, principal component analysis of the spots was performed using the *RunPCA* function. Uniform Manifold Approximation and Projection (UMAP) layouts and nearest-neighbor graphs were generated using the top 25 components. Different resolutions for spatial clustering were explored to determine the optimal resolution for analyzing tissue structure. Each sample’s data were stored as Seurat objects.

### Spatial spot deconvolution of BM with scRNA-seq reference dataset

Single-cell RNA sequencing (scRNA-seq) Seurat objects, predefined for cell types and stored as RDS files, were loaded using the *readRDS* function. Cells from 9 healthy BM samples (n = 62,862 cells) and the AML cells subset from 7 patients with diploid karyotype BM (n = 16,167 cells) were integrated using the merge function. The combined dataset was then normalized with *SCTransform*, and the top 2,500 features were identified using *FindVariableFeatures*. A total of 79,029 cells were scaled and centered using the *ScaleData* function. Dimensional reduction was performed on the scaled data with principal component analysis (PCA) using the *RunPCA* function, selecting 50 principal components with assigning npcs parameter as 50. Batch effects potentially arising from this integration were removed using the *RunHarmony* function from the Harmony package.^38^ Neighborhoods and clusters were computed using the top 30 principal components, followed by calculation of UMAP layouts.

To deconvolve cell types in spatial spots utilizing predefined cell-type labels from the reference scRNA-seq dataset, transfer learning was employed using the *Seurat* package. Integration anchors between the ST data and the scRNA-seq reference were established using the *FindTransferAnchors* function, employing SCT normalization. During this alignment, residuals were not recomputed, as specified by setting the *recompute.residuals* parameter to False.

Following anchor identification, cell-type annotations were transferred from the scRNA-seq dataset to the ST data using the *TransferData* function. This function applies a weighted nearest neighbor approach, leveraging weights derived from the first 25 principal components of the PCA embeddings specific to the ST data. The annotations transferred in this manner were stored in a new assay within the ST dataset.

To further categorize the spatial spots, deconvolution scores from the assigned cell-types were used. Spots with AML prediction scores above the median value (median = 0.15) were classified as AML-enriched, while those below this threshold were designated as AML-depleted. For detailed deconvolution AML-enriched spots were subsetted. Only AML cells from our scRNA-seq data, along with predefined AML cell state information, were used to deconvolve AML cell populations within the AML-enriched spots using the same transfer learning approach.

### Spatial spot deconvolution of EM samples

EM sample data stored as Seurat objects were converted to *SpaCET* objects using the *convert.Seurat* function from the *SpaCET* package (version 1.1.0).^39^ The *SpaCET.deconvolution* function was then used to deconvolve these data using the Pan Cancer dictionary, which includes average all cancer type-specific expression signatures for 30 solid tumors, with the *cancerType* parameter set to ‘PANCAN’. To identify malignant states within our tissues, the *SpaCET.deconvolution.malignant* function was used. The deconvolution results were added to the EM samples’ Seurat object as a new assay using the *addTo.Seurat* function. Subsequently, these scores were visualized and assessed using the *SpatialFeaturePlot* in Seurat package.

In the EM1 sample, the leukemic population within the cluster 1 was isolated using the *subset* function. Subsequently, similar to our approach with the BM1 sample, the AML cell population and their corresponding cell state information present in our scRNA-seq dataset were specifically deconvolved using a transfer learning approach.

### Spatial Co-Localization Analysis

Deconvolution scores for leukemic populations were used to define spots with high and low leukemic scores in BM. Deconvolution scores that contain all cellular composition information were stored as a new assay in the BM2’ Seurat object. Wilcoxon rank sum test applied to define differential scores of cell types with the *RunDEtest* function from the SCP (version 0.5.1) package. Results were visualized with the VolcanoPlot function.

In EM, deconvolution scores were extracted from the Seurat object. The scores were then transposed, and a Pearson correlation matrix was calculated to evaluate the pairwise correlation between cell type abundances. Mean deconvolution was computed to determine the abundance of each cell type.

### Spatial Cell-Cell Communication Analysis

Cell labels were assigned to spots based on prediction probabilities from a custom function *assignLabels*. The data was prepared for cell-cell communication analysis using the CellChat package^40^ (version 2.1.2), with spatial locations and conversion factors obtained from the 10X Visium data. A CellChat object was created with *createCellChat* function, and ligand-receptor interaction databases were set and subsetted using *subsetDB* to focus on secreted signaling. Overexpressed genes and interactions were identified with *identifyOverExpressedGenes* and *identifyOverExpressedInteractions*.

Communication probabilities were computed using computeCommunProb, and the cell-cell communication network was inferred. Pathway analysis was performed with *computeCommunProbPathway*, *netAnalysis_computeCentrality*, and *netAnalysis_signalingRole_network*. Specific pathways such as CXCL highlighted using *netVisual_* functions and *plotGeneExpression*.

### SpatialTime and Distance Analysis

SpatialTime analysis was performed following established methods.^41,42^ Initially, contours were manually drawn on Visium CytAssist image of trabecula with ImageJ2 (version 2.14.0). For each spatial spot in the BM, distances to the closest contoured surface were computed and scaled from 0 (adjacent to the surface) to 1 (furthest from the surface). The median of the calculated SpatialTime values was used to define proximity and distality: spots above this calculations median value (median = 0.15) were classified as distal, while those below it were classified as proximal. Deconvolution results were visualized using the *FeatureStatPlot* function of the *R* package *SCP* to display the concentration of features at different distances relative to the trabeculae.

### Pathway Analysis and Inflammation Classification

Pathway analysis was conducted using curated gene sets (hallmark), obtained from the Molecular Signatures Database (MSigDB). These gene sets were individually scored for each sample using the AUCell^43^ (version 1.24.0) pipeline, and the resulting scores were integrated into our datasets.

Spatial coordinates and pathway scores for inflammatory pathways were extracted from the Seurat objects including inflammatory response, IL6/JAK/STAT3 signaling, IFNγ Response, IFNα Response, TNFα/NF-κB signaling, complement, and IL2/STAT5 signaling. Pathway scores were normalized to a 0-1 scale, and a composite score was calculated as the mean of these normalized scores. Coordinates and composite scores were combined into a data frame, and Jenks Natural Breaks method was applied to categorize composite scores into four categories: Low, Medium-Low, Medium-High, and High. A spatial map was created to visualize composite inflammatory pathway activity. Box plots were generated to compare composite scores across related classification (Clusters, CXCL12-CXCR4, PI3K/AKT/mTOR, Trans-differentiation). Composite scores and classes were added to the Seurat object’s metadata, ensuring alignment with the spots in the object. Spatial feature and dimension plots were generated to visualize composite scores and categories within the spatial context of the tissue sections.

### Opal Multiplexed Fluorescent Immunohistochemistry

Near-adjacent tissue sections from PT1 and PT2 were baked overnight, deparaffinized and quenched using hydrogen peroxide and 470 nm light. Slides underwent heat-induced epitope retrieval, followed by blocking and incubation with primary antibodies (Table S5). Horseradish peroxidase and Opal fluorophores were applied, with antibodies stripped between each round of Opal staining to enable multiplexing. DAPI (4’,6-diamidino-2-phenylindole) was used as a nuclear counterstain. All procedures were automated on the NanoVIP 100 stainer (Biogenex), and slides were imaged at 0.25 µm resolution using the PhenoImager HT 2.0 (Akoya Biosciences), with unmixing performed at acquisition using custom fluorescence libraries.

### Image Analysis and Alignment of Opal Validation Images

Unmixed Opal images were aligned with Visium Cytassist reference images by applying 15-25 common anchor points within the TissueAlign^TM^ module of Visiopharm software (version 2024.07.1.16912 x64). Opal images were then segmented using a combination of machine-learning-based algorithms and manual adjustments to remove artifacts, bone, and empty space, defining a single region of interest for each tissue. This region was further segmented into cells based on DAPI channel using a deep learning (U-net) approach. Cell intensities first had a 3x3 median filter applied as a noise reduction filter. For each marker, mean cell intensity was calculated based on the pixels comprising 70-95% of each cell’s intensity to account for nuclei and exclude outliers. The mean was then arcsinh-transformed to normalize the distribution. Cells were assigned to corresponding Visium spots based on the intersection of their coordinates in the reference image with the boundaries of Visium spots, and mean marker intensity was calculated for each spot.

## Results

### Comparative spatial transcriptomics analysis of BM and EM tissues in AML

Trephine BM biopsies from two AML patients, along with a cutaneous myeloid sarcoma (skin) punch biopsy (EM1) from patient 1 (PT1) and a surgical biopsy of a mediastinal lymph node EM mass (EM2) from patient 2 (PT2), were collected at the time of AML diagnosis (pre-treatment) and were used for Visium-based ST profiling (**Figure 1A**). Clinical characteristics of these patients are summarized in **Table S1**. Briefly, PT1 (age 41 years; male) developed cutaneous myeloid sarcoma with concomitant medullary (BM) leukemia, characterized by 30% myeloblasts in the BM (BM1). PT2 (age 83 years; male) presented with a mediastinal mass but showed no histopathologic confirmation of medullary involvement of his AML (2% myeloblasts detected in BM2) (**Figure S1A-C**). While the targeted mutation panel in EM2 was negative for mutations, the BM of PT1 revealed mutations *in NPM1, DNMT3A, IDH1, IDH2, FLT3, SF3B1,* and *KRAS*. Both patients’ BM samples had diploid cytogenetics.

**Figure 1:**
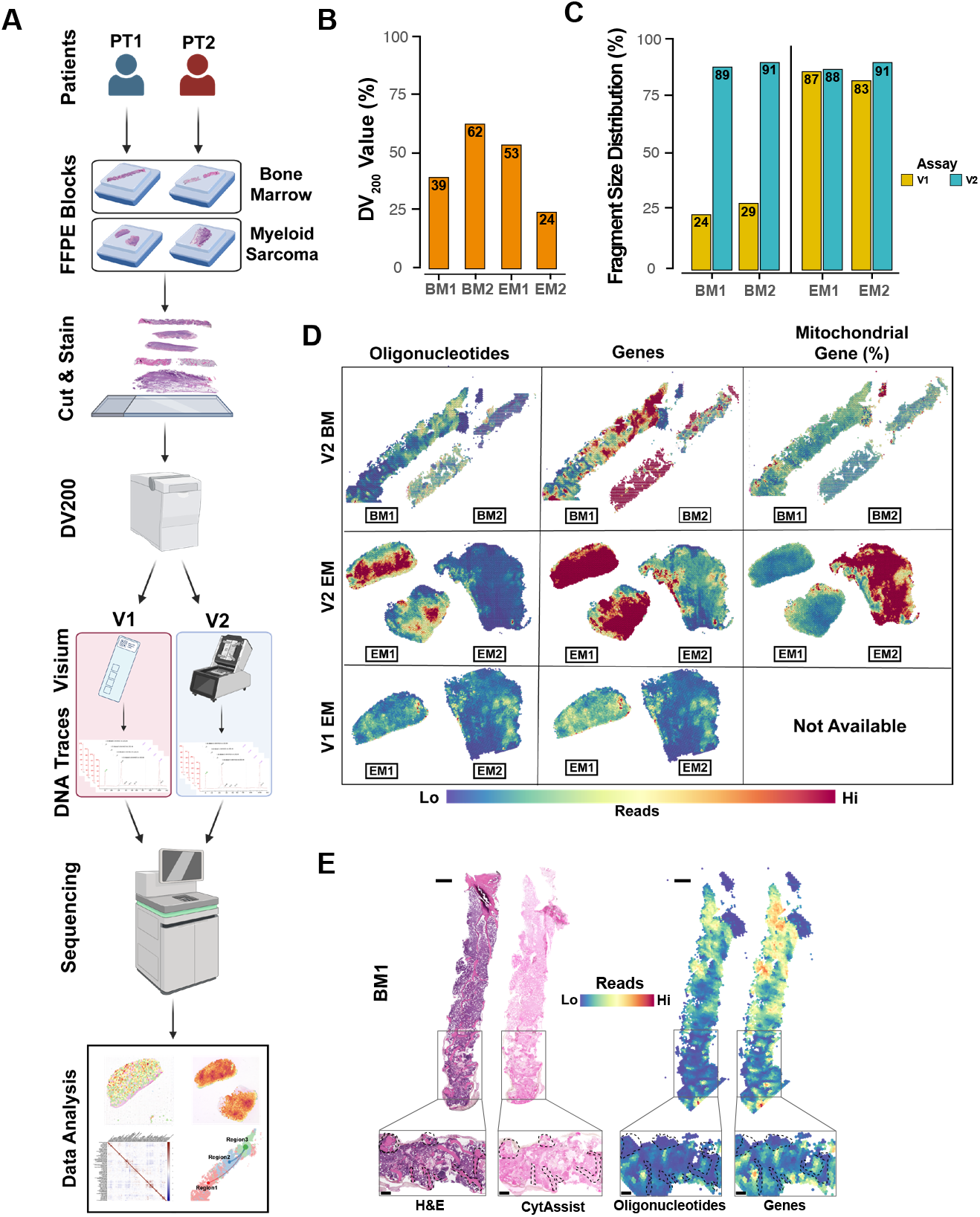
Study overview and Quality control comparison of Visium v1 and Visium v2. **(A)** Schematic representation of the study workflow, visualized by BioRender (https://biorender.com/). Concurrent Bone Marrow (BM) and Extramedullary (EM) (EM1 from skin and EM2 from lymph node) formalin-fixed paraffin-embedded samples from 2 newly diagnosed patients with acute myeloid leukemia (AML) (PT1 and PT2) were obtained. Each sample was sectioned and stained with hematoxylin and eosin (H&E). RNA integrity was evaluated using DV_200_ measurements before library preparation. Visium ST assays v1 and CytAssist v2 were performed on each sample. After library preparation, cDNA traces were assessed, and samples meeting the criteria were sequenced for further analysis. **(B)** DV_200_ values for BM1, BM2, EM1, and EM2 tissues, indicating RNA integrity before library preparation **(C)** Fragment size distribution percentages calculated within the 200-1000 bp range for DNA libraries prepared using v1 and v2 assays. BM samples showed significant improvement with v2 compared to v1. EM samples were within the acceptable range for both assays. **(D)** Spatial mapping by Seurat of captured oligonucleotides, genes, and mitochondrial gene percentages in BM and EM tissues. The v2 assay demonstrated higher oligonucleotide and gene capturing compared to v1, with mitochondrial gene percentages available only for v2. **(E)** Histological overlay of H&E-stained BM1 section with CytAssist image and spatial transcriptomics data, The image shows preserved structural integrity of bone regions, allowing detailed spatial analysis. The main tissue scale bar indicating 1 mm, while the zoomed-in panels, highlighting the boxed regions, denoting 400 µm.

RNA fragmentation before Visium library preparation (pre-library), measured as the percentage of fragments > 200 nucleotides (DV_200_), varied across the 4 analyzed samples (range 24-62%; mean DV_200_ = 44.5, standard deviation = 16.6) (**Figure 1B**, **Figure S1D**). The EM2 sample showed the lowest pre-library DV_200_ value at 24%, whereas the BM2 sample from the same patient displayed the highest pre-library DV_200_ metric at 62%. Then, to compare the assays and prepare the libraries for ST profiling, we used two Visium-based assays; version 1 (v1) (direct placement of each tissue section on a spatially-barcoded Visium slide), and version 2 (v2) (utilizes a tissue transfer-based approach in the CytAssist instrument) on all 4 patient samples in parallel, totaling 8 libraries. For BM tissues, post-library DNA traces for v1 libraries were markedly lower compared to v2 (24% and 29% for v1 versus 89% and 91% for BM1 and BM2, respectively) (**Figure 1C**, **Figure S1E, F**), precluding further sequencing of v1 BM libraries. However, for EM tissues, the post-library DNA tracing was similar between v1 and v2 (EM1: 87% and EM2: 83% for v1, and EM1: 88% and EM2: 91% for v2). This suggests that Visium v2 automated tissue transfer provides a more reliable performance compared to v1, even with low pre-library DV_200_ measures.

Spatial mapping of oligonucleotide and gene counts obtained by SpaceRanger pipeline revealed that compared to v1, v2 assay had more oligonucleotides detected (mean nCount; 1292.76 for v1 vs 11072.37 for v2) and genes identified (mean nFeature; 955.19 for v1 vs 4735.67 for v2) per sample (**Figure 1D**; **Figures S1G-I**). As expected, the median oligonucleotides detected and genes identified were positively correlated (r = 0.996, p = 1.4 x 10-5) (**Figure S1J**). Mitochondrial gene percentages, available for v2 only, were highest in EM2, consistent with a necrotic phenotype on the histopathologic assessment (**Figure S1B**). Of note, prelib DV_200_ values were negatively correlated with the median percentage of mitochondrial genes detected (r = -0.838; p = 0.1625) (**Figure S1L**). These mitochondrial reads were higher at the edge of the samples (**Figure S1K**). To assess whether the structural integrity of bone regions was maintained during tissue processing, we overlaid the hematoxylin and eosin (H&E) image generated on the same Visium slide of BM tissues with the ST data derived from SpaceRanger. Indeed, spots in bone trabeculae areas that usually have low cell abundance and tendency to come off during tissue processing were still maintained (**Figure 1E**, **Figure S1M**), allowing downstream analysis of spatial BM components using the CytAssist transfer-based approach.

### Spatial clustering and spot deconvolution reveal tissue morphology

To our knowledge, Visium-based approaches in AML is not explored. This is particularly pertinent due to decalcification, RNA quality and unique bone-rich environment of AML compared to solid cancers. We thus adapted the median absolute deviation (MAD) on the spot level quality metrics to filter spots based on the normal distribution of mitochondrial content and expression profile (**Methods, Figure S2A-F**). To spatially delineate spot-level cellular information, we then deconvolved individual spots using probabilistic label transfer workflow^44^ based on our in-house generated scRNA BM reference (**Figure 2A**). This reference map consisted of 79,029 cells collected from 9 healthy BM donors and 7 patients with AML with diploid cytogenetics including both newly generated scRNA data and previous works.^13,34^ The reference map was clustered into a total of 21 different cell types, including T cells (CD4^+^ and CD8^+^ naïve, effector, and memory T cells, T regulatory [Treg] cells, and unconventional T cells), other immune cells (Natural killer [NK] cells, B cells and plasma cells), hematopoietic progenitors (Hematopoietic stem cells [HSCs], common lymphoid progenitors [CLPs], granulocyte-monocyte progenitors [GMPs]), myeloid cells (megakaryocytes/platelets, monocytes, early and late erythroid cells, conventional and plasmacytoid dendritic cells) and leukemic (AML) cell populations. The identities of these cell types were verified using canonical gene expression, as previously done.^13,45–47^

**Figure 2:**
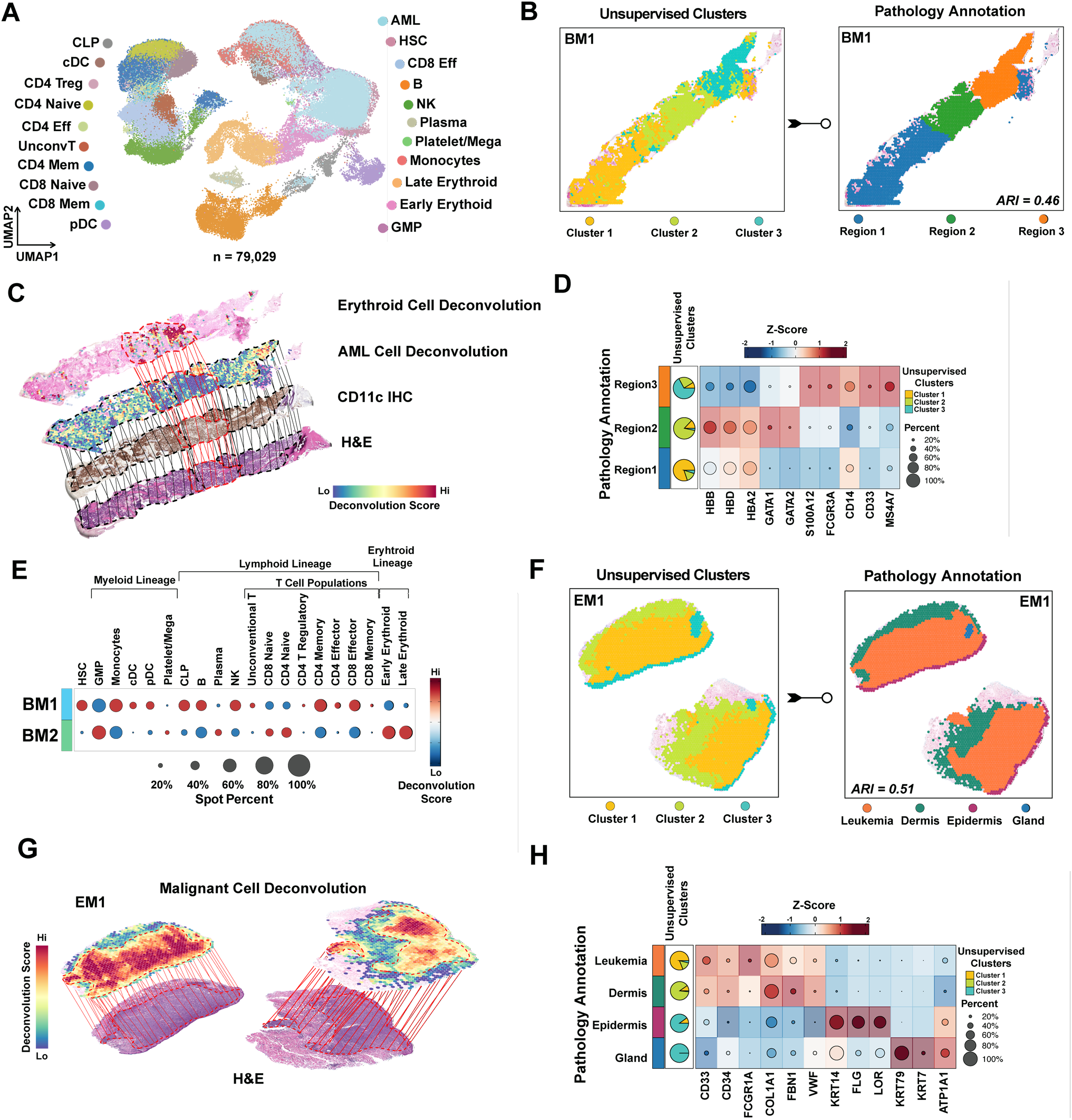
Spatial clustering and cell deconvolution in AML tissues. **(A)** Uniform Manifold Approximation and Projection (UMAP) plot showing 21 different cell types identified from the scRNA reference map, derived from 79,029 cells from 9 healthy donors and 7 diploid AML patients. AML, acute myeloid leukemia; cDC, classical dendritic cells; CLP, common lymphoid progenitors; Eff, effector T cells; GMP, granulocyte-monocyte progenitors; HSC, hematopoietic stem cells; Mega, megakaryocytes; Mem, memory T cells; NK, natural killer cells; pDC, plasmacytoid dendritic cells; Treg, T regulatory cells; UnconvT, unconventional T cells. **(B)** Unsupervised clustering and pathology annotation for BM1 projected spatial map, revealing 3 distinct regions with an adjusted Rand index (ARI) of 0.46. **(C)** Spatial deconvolution of BM1 tissue, showing erythroid and AML cell populations, with CD11c immunohistochemistry (IHC) overlaid on an H&E-stained image. Dotted red lines represent regions enriched for the erythroid cell population. Dotted black lines indicate regions enriched for the AML cell population. Solid lines represent regions that overlapped with other tissue sections. **(D)** Heatmap showing canonical marker expression in pathology annotations, with matching unsupervised cluster distribution represented as a pie chart. **(E)** Dot plot depicting the percentage and deconvolution score of each cell type in BM1 versus BM2 tissues. **(F)** Spatial map of unsupervised clusters and pathology annotation for EM1, with an ARI of 0.51, identifying distinct tissue regions. **(G)** Spatial deconvolution of EM1 tissue obtained from SpaCET algorithm, showing malignant cell distribution overlaid on the H&E image. **(H)** Heatmap of canonical marker expression in EM1 regions, validating transcriptional segregation and matching pathologist-defined regions. Markers for leukemic population and dermis regions show shared expression profiles. Unsupervised cluster overlap is represented as pie charts, with pathology annotation.

We next performed shared nearest neighbor (SNN) modularity optimization^48^ based unsupervised clustering and compared the cluster profiling to hematopathologist-identified histopathologic tissue annotation i.e. ground truth (**Figure 2B**, **Figure S3B-D**). For instance, in BM1, unsupervised clustering segmented the BM into 3 distinct clusters. Independently, the pathologist also identified 3 distinct regions with an adjusted rand index (ARI) of 0.46 (demonstrating moderate overlap). Further, regions 1, 2 and 3 were annotated by the pathologist as mixed, erythroid-enriched and monocytic/leukemia-enriched, respectively, which was consistent with the deconvolution analysis (**Figure 2C**, **Figure S3E**). To further validate the deconvolution-derived spot annotation, images stained with H&E, CD11c, CD3 and myeloperoxidase (MPO) on serial slides imaged, overlaid with the Visium slides and were evaluated by a hematopathologist (**Figure S3A,F).** This was further supported by differential expressed gene analysis and canonical markers on ST data of these regions by erythroid cells (*HBB, HBA2, GATA1, GATA2*), monocytes (*S100A12, FCGR3A, CD14, MS4A7*), and early myeloid cells (*CD33*) (**Figure 2D**, **Table S2**). We detected all 21/21 (100%) of the cell types in our reference map. In BM1, the most abundant cell type was AML with late erythroid cells and monocytes, whereas in BM2 it was late erythroid cells with GMPs were most common (**Figure S3G**). Compared to BM2 (no leukemia detected), BM1 (∼30% leukemic blasts) had slightly higher prevalence of effector and memory T-cell populations, while BM2 had high abundance of erythroid-lineage populations (**Figure 2E**).

Since EM tissues are non-medullary tissues and contains tissues such as epidermis, dermis, germinal centers, glands, etc., we applied the Spatial Cellular Estimator for Tumors (*SpaCET*) algorithm^39^ optimized for solid non-BM cancers to identify tumor regions (**Methods**). Unsupervised clustering segmented the tissue into 3 distinct regions, which mostly overlapped with histopathological annotations except that of glandular tissue (*ARI = 0.51*) (**Figure 2F**). Similar to our approach in BM samples, deconvolution results were confirmed with the histopathologic annotations using the same-slide H&E digital images (**Figure 2G**, **Figure S3A**). In EM1, macrophages were the predominant cell population, whereas cancer-associated fibroblasts and endothelial cells were the most enriched in EM2 (**Figure S3H**). Tissue-specific markers and differential expressed genes validated transcriptional segregation, revealing dermis infiltration by leukemic cells in EM1, and confirmed the consistency of unsupervised clusters with pathologist-defined regions (**Figure 2H**, **Figure S3I, Table S3**). These results support our use of deconvolution for spot-level annotation allowing further downstream analysis.

### AML-population focused analysis demonstrates spatial heterogeneity by cell compositions in both BM and EM tissues

Using the median deconvolution score, we assigned spots as high leukemic scores (HLS) (greater than the median of 0.15) or low leukemic scores (LLS) (**Figure 3A**). In BM1, cluster-based zonation revealed that HLS spots were ∼14% more abundant in cluster 3, while LLS spots exhibited a 20% increase in cluster 2 (**Figure 3B**). GMPs and monocytes were both localized in HLS spots, with monocytes predominantly in cluster 3 and GMPs scattered across clusters (**Figure 3C**, **D**). CD8^+^ naive cells were scattered throughout, with a higher concentration in cluster 1, which contained mixed cell populations (**Figure 3E**). Spots harboring late erythroid cells were predominantly found in the cluster 2 where HLS spots were least abundant (22.9% of HLS spots) and encompassed the most LLS spots (42.9% of LLS spots). In EM1, immune cell populations were found to be localized together within the tissue. Notably, macrophages, which had the highest abundance in EM1, showed strong co-localization with classical dendritic cells (*Pearson correlation coefficient (r) = 0.67)* and significant associations with cancer-associated fibroblasts (*r* = 0.37) (**Figure 3F**). Spatial mapping revealed that macrophages and classical dendritic cells co-localized in regions with tumor infiltration in the dermis (**Figure 3G**). These results revealed that GMPs and monocytes co-localize with leukemic-enriched populations in the BM, while macrophages are concentrated in tumor-infiltrated regions in the EM.

**Figure 3:**
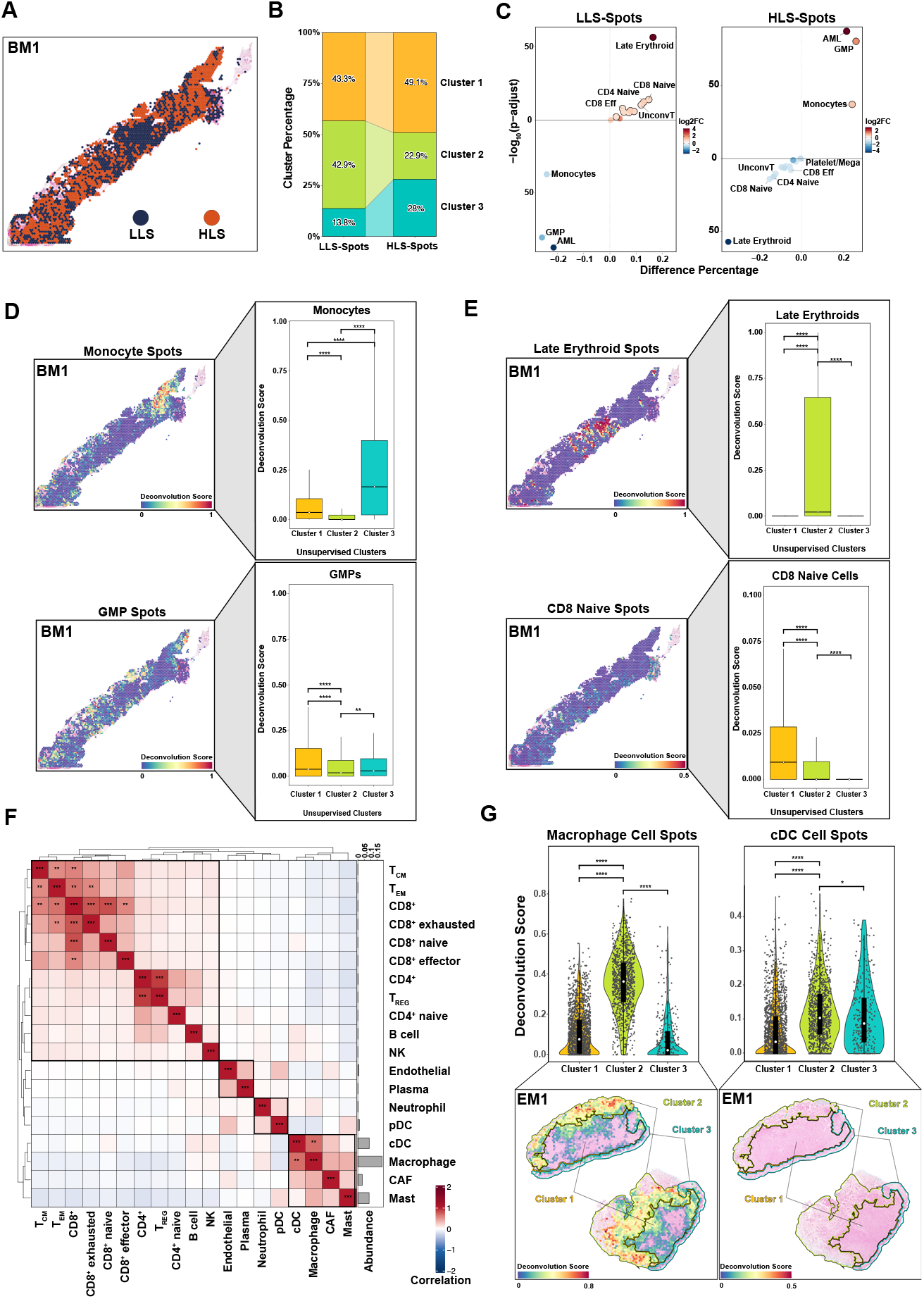
Spatial heterogeneity of AML populations in BM and EM tissues. **(A)** Spatial map of BM1 showing spots with high leukemic scores (HLS) (above median AML cells deconvolution score; >0.15) (orange) and low leukemic scores (LLS) (below median AML cells deconvolution score; <=0.15) (darkblue). **(B)** Stacked bar plot shows cluster-based distribution of HLS and LLS spots. **(C)** Volcano plot shows differential co-localization of cell populations within HLS and LLS spots. Deconvolution scores were compared by Wilcoxon rank sum test. **(D)** Spatial deconvolution and distribution (represented as a box plot) of monocytes and GMPs in HLS spots in BM1, with deconvolution scores indicating higher abundance in cluster 3. Median values are shown as white dots on a black line. **(E)** Spatial deconvolution and distribution of late erythroid cells and CD8 naïve cells in BM1, showing their localization in LLS spots. **(F)** Correlation heatmap of cell populations in EM1, highlighting significant co-localization between macrophages and classical dendritic cells (cDCs). Pearson absolute correlation: ***>0.7, **>0.5. **(G)** Spatial mapping of macrophage and cDC spots in EM1, showing their co- localization in tumor-infiltrated dermis clusterd (clusters 1-2). *p < 0.05, **p < 0.01, ***p < 0.001, ****p < 0.0001, Wilcoxon rank sum test.

### Inferred pathway and cell communication analysis uncover interaction of CXCL12-CXCR4 axis with inflammation

Differential gene expression analysis was conducted on HLS spots relative to LLS spots in BM1 (**Figure S4A, TableS4**). We discovered that genes highly expressed in HLS-spots were concentrated in cluster 3 (**Figure S4B**). Among these genes associated with leukemic regions in the BM, those such as *CD70*,^49^ *TMEM176B*,^50,51^ *TP53INP2*,^52^ and *TNFSF13B*,^53^ which are associated with immune regulation and tumor progression, were also found to be expressed in the leukemic regions of the same patient’s EM sample (**Figure 4A**).

**Figure 4:**
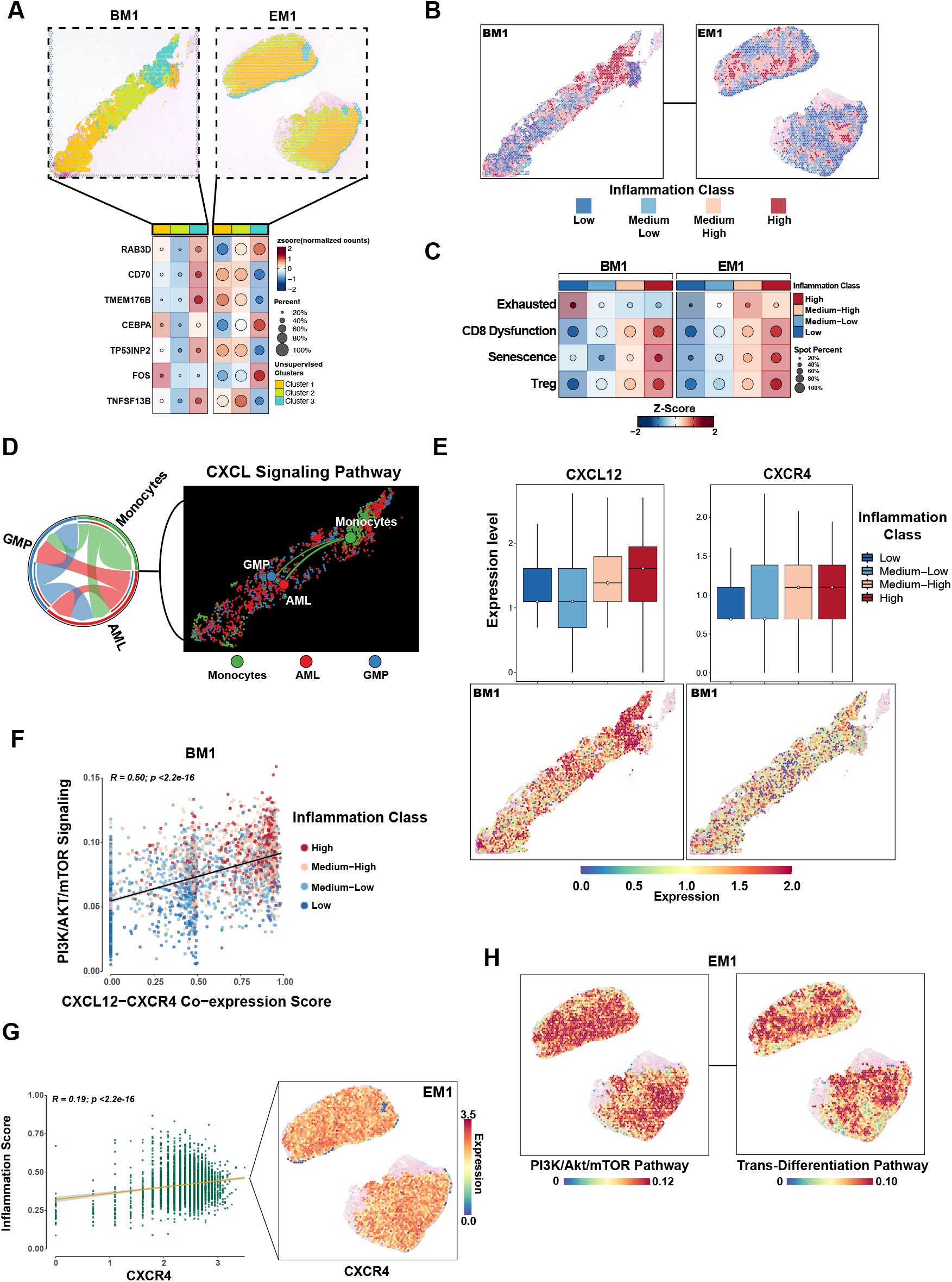
Interaction of the CXCL12-CXCR4 axis with inflammation in AML tissues. **(A)** The heatmap shows differential gene expression analysis of HLS spots relative to LLS spots in BM1, highlighting genes associated with immune regulation and tumor progression. Genes highly expressed in HLS spots of BM1, particularly in cluster 3, show a similar expression profile to that of the leukemic-enriched cluster 1 of EM1. **(B)** Spatial inflammation class distribution in BM1 and EM1, based on composite inflammation scores from inflammation-related hallmark pathways, shows medium-high and high inflammation spots concentrated in leukemic-enriched regions (cluster 3 in BM1 and cluster 1 in EM1). Classes were defined using Jenks natural breaks optimization. **(C)** Dot plot shows T cell subtypes (exhausted, dysfunction, senescence, regulatory) localization based on inflammation class on PT1. **(D)** Spatial and circus plot of AML- GMP-monocyte interactions and interactions strength among spots predicted by CellChat. **(E)** Box plots showing expression levels of CXCL12 and CXCR4 in BM1, stratified by inflammation class. Spatial maps indicate expression distribution. **(F)** Correlation between PI3K/AKT/mTOR signaling and CXCL12-CXCR4 co-expression scores in BM1, stratified by inflammation class. T-distribution was used to evaluate the significance of Pearson correlation. **(G)** Relationship between CXCR4 expression and inflammation score in EM1, with spatial maps showing expression distribution. **(H)** Spatial mapping of PI3K/AKT/mTOR and trans-differentiation pathway (epithelial-mesenchymal transition from hallmark pathways) in EM1.

Interestingly, pathway profiling revealed the concomitant BM and EM samples of same patients also displayed similar molecular signatures (**Figure S4C**), suggesting similar biologic programs governing medullary and EM leukemia. For instance, in BM1, pathways related to inflammation (IFNα, IFNγ, inflammatory response, and TGF-β), and energy metabolism (oxidative-phosphorylation [OXPHOS] and glycolysis), were upregulated in both sample of PT1 but were slightly downregulated in the samples from PT2. Additionally, epithelial-mesenchymal transition (EMT)-like signatures linked to neoplastic cell migration and trans-differentiation were prominent in our EM samples. Comparing pathway activities between spatial clusters within each tissue, we found a similar profiles between cluster 3 of BM1 and the leukemic population cluster of EM1 (**Figure S4D**).

Dysregulated inflammatory pathways in the BM microenvironment contribute to leukemogenesis and leukemic blast maintenance in AML.^54,55^ To identify the inflammation niche in our AML-affected BM sample (BM1), we defined a composite inflammation score using inflammation-related hallmark pathways (inflammatory response, IL6/JAK/STAT3 signaling, IFNα and IFNγ, response, TNFα signaling via NF-κB, complement, and IL2/STAT5 signaling) and then clustered the spatial data using Jenks natural breaks optimization (**Methods**, **Figure 4B**). We observed a high inflammation profile in BM1 that was concentrated in cluster 3, mirroring the high inflammation scores found in the leukemic population of EM1 (**Figure S4E,F**). Further, we found that the hypoxic environment (Hypoxia, reactive oxygen species and HIF1A pathways) intensified within the highly inflammatory niche (**Figure S4G**). Taken together, our findings demonstrate that the leukemic population in distinct tissues from the same patient can exhibit similar biologic profiles.

Deeper characterization of T cells revealed that their exhausted profiles, despite being in low abundance, are observed in low-inflammatory regions within BM1 but in high-inflammatory regions at extramedullary site. Additionally, CD8 dysfunction^45^, senescence^47^, and Treg^56^ profiles are associated with high-inflammatory regions in both tissues (**Figure 4C**, **Figure S4H**). We then evaluated the spatial cell-cell interactions.^40^ Based on deconvolution results; each spot was labeled according to the cell type with the maximum prediction probability (**Figure S5A**). Our analysis revealed that CXCL pathways between annotated spots exhibited the highest signaling strength (**Figure S5B**). Specifically, the CXCL12-CXCR4 axis showed that monocytes and GMPs co-localized with AML cells in HLS spots (**Figure 3C-D**), indicating communication through this pathway (**Figure 4D**, **Figure S5C-D**). Examining the relationship of the CXCL12- CXCR4 pair within the inflammatory niche, we found that both the CXCL12 ligand and the CXCR4 receptor showed high expression levels in medium-high and high inflammation spots in BM1 (**Figure 4E**).

The PI3K/AKT/mTOR pathway, a downstream target of these chemokines, was highly correlated with CXCL12-CXCR4 co-expression in regions of elevated inflammation (**Figure 4F**, **Figure S5E,F**).^57^ This pathway is also involved in the direct induction of trans-differentiation (**Figure S5G**),^58^ aligning with our findings of elevated EMT pathway in these regions. To understand the involvement of this PI3K/AKT/mTOR through CXCL12-CXCR4 pathway at the EM site, we examined the spatial expression profiles of its ligand-receptor communication. We observed that CXCR4 was abundantly expressed throughout the tissue and correlated with the composite inflammation score (**Figure 4G**). Additionally, the PI3K/AKT/mTOR pathway and trans-differentiation states were active in regions with high inflammation in EM (**Figure 4H**, **Figure S5H,I**). These observations suggest that leukemic populations in the BM concentrate within the inflammatory niche via CXCL12-CXCR4 chemokine signaling, activating trans- differentiation through the PI3K/AKT/mTOR pathway. Additionally, the elevated levels of CXCR4 and inflammation in EM suggest a possible path for medullary leukemia to migrate EM tissues.

### Multiplexed immunohistochemistry confirms spatial patterns observed in ST data

To validate our ST findings, we performed mfIHC on near-adjacent tissue sections, obtaining spatial proteomic information at single-cell resolution. These data were aligned with the corresponding Visium samples to create spot-level information (**Figure 5A**, **Figure S6A**). Using phenotypic markers—CD33 (leukemic cells), CD68 (monocytic phenotype), and CD71 (erythroid phenotype)—we validated our annotation approach, observing similar spatial patterns between mfIHC and ST data (**Figure 5B**, **Figure S6B,D-E**). The cluster-based distribution of protein expression mirrored the ST data, with cluster 3 enriched for leukemic and monocytic populations and cluster 2 enriched for erythroid cells in BM1 (**Figure 5C**).

**Figure 5:**
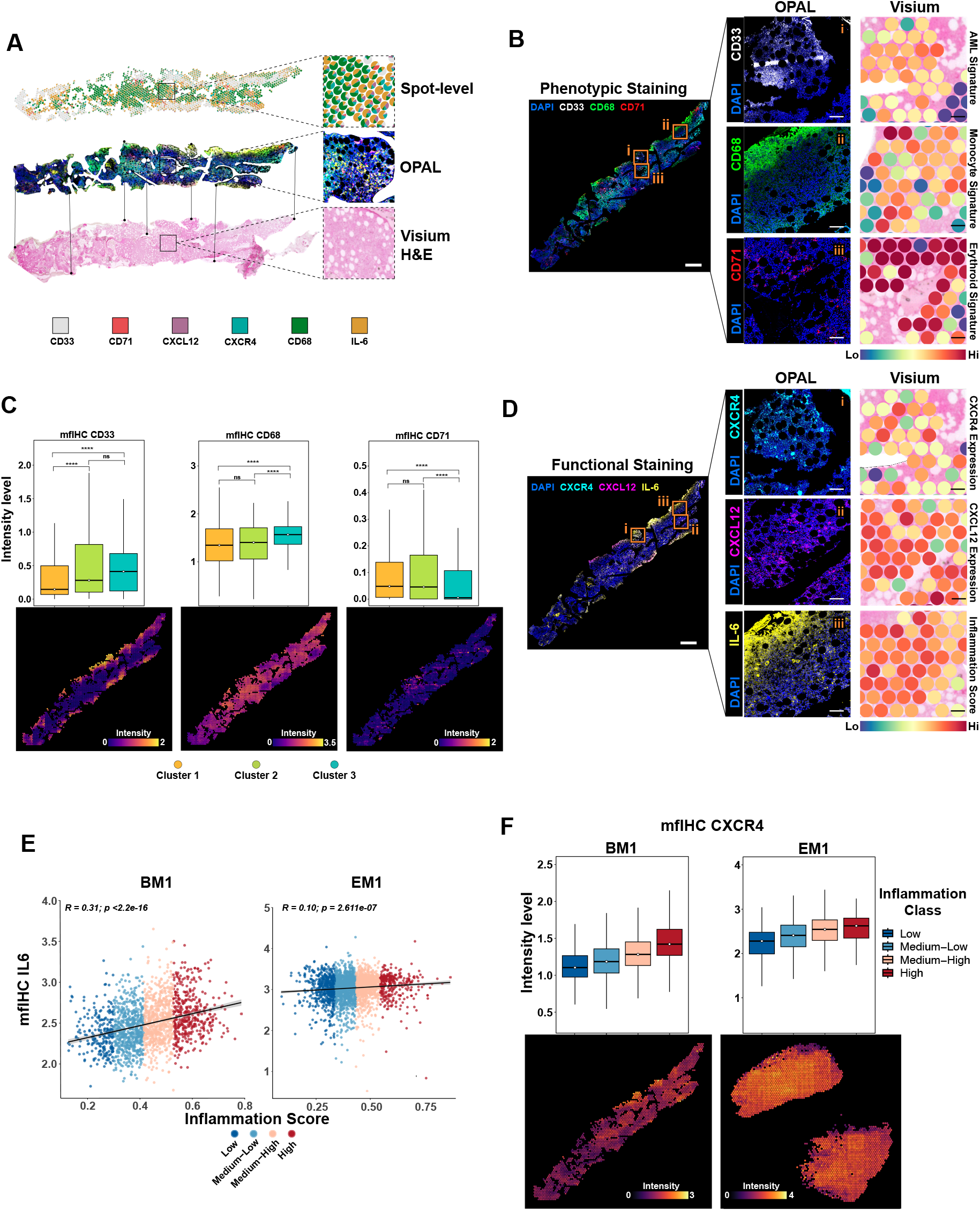
Validation of ST findings with multiplexed fluorescence immunohistochemistry. **(A)** Spatial map illustrating the alignment of Visium and OPAL-stained slides to obtain spot-level proteomic information at single-cell resolution. Dots and lines between OPAL and Visium H&E are representative to using common anchor point. **(B)** OPAL-stained images with phenotypic markers (CD33, CD68, CD71) and corresponding Visium spots exhibit similar spatial patterns. The top panel shows whole-slide pseudo-clor images with ROI squares, while left and right panels display zoomed OPAL and Visium images from identical locations. **(C)** Box and spatial plots of mfIHC intensities for phenotypic markers across ST-defined clusters, highlighting enrichment of leukemic and monocytic populations in cluster 3 and erythroid populations in cluster 2 at BM1. **(D)** OPAL and Visium images for functional markers confirm alignment of protein expression with ST data (**E**) Pearson correlation between IL-6 mfIHC intensities and the composite inflammation score demonstrates a positive association, with higher IL-6 levels observed in high-inflammation regions at PT1 samples (**F**) Box and spatial plots indicate increased CXCR4 intensity withing high-inflammatory niches in BM1, corroborating ST findings. Scale bars: 1 mm (whole-slide panels) and 100 µm (selected region panels).

We further validated our ST signatures using functional markers; CXCL12-CXCR4 (chemokine and receptor axis for cell migration), and IL-6 (inflammation marker). Histological validation showed consistent localization of these markers with the ST data (**Figure 5D**, **Figure S6C-E**). IL-6 mfIHC intensities correlated with our composite inflammation score, confirming accurate spatial assignment of the inflammatory niche (**Figure 5E**, **Figure S6F**). Additionally, CXCR4 protein expression was enriched in highly inflammatory regions in PT1 samples (**Figure 5F**). CXCL12 expression was elevated in inflammatory regions in BM1 but was region independent in EM1 as we observed in ST data (**Figure S6G**). Taken together, the orthogonal approach we applied validates the accuracy of our ST approach and underscores the importance of combining spatial multi-omics techniques in such studies.

### Deconvolution of leukemic-enriched spots reveals the localization of different differentiation states of AML cells within inflammatory and endosteal niches

We next applied linear mixed model annotation^59^ to classify AML cells (n=16,167) from scRNA into their differentiation states relative to the hierarchies in healthy BM controls (n=20,778 cells).^60^ This classification defined AML cells as: primitive-like (combination of HSC-like, common myeloid progenitor/lymphoid-primed multipotent progenitor-like; n= 5,039), GMP-like (n=6,432), erythroid-like (n= 1,816), lymphoid-like (n = 55), and committed-like (combination of monocyte-like, basophil-like, dendritic cell-like; n=2,825) (**Figure 6A**). We then applied this classification to BM HLS spots (n=1,271 spots) and EM1 leukemic-enriched spots (n = 1,726 spots) (**Figure 6B**, **C**) to obtain a spot-level classification of hierarchies. Cluster-based niches revealed that committed-like populations were located distally to the primitive-like populations (**Figure S7B, C**). Deconvolution scores indicated a lower abundance of primitive-like cells compared to committed-like cells in EM tissue (**Figure S7D,E**). We found that committed-like AML populations were concentrated in inflammatory niches in both BM and EM tissues (**Figure 6D,E**). We then applied the SpatialTime pipeline^41^(**Methods**) to measure the spatial localization of HLS spots relative to the trabecular bone regions and based on the degree of AML differentiation (**Figure 6F**, **Figure S7F**). We found that primitive-like cell populations were localized proximally to the bone, while GMP- and committed-like populations were localized distally (**Figure 6G**). To validate these findings, we performed a GeoMx-based whole-transcriptome microdissection-based assay also known as digital spatial profiling (DSP) on 13 BM regions from 3 independent newly diagnosed AML patients (mean age: 73 years; 3 male). (**Figure S7G-H**). CD34 and CD68 protein markers were used to define the leukemic regions covering primitive and more differentiated cells (**Figure S7I-J**). Congruent with our Visium ST analysis, phenotypically primitive-like cells were detected proximal to the bone, while more differentiated cell states were predominantly found in regions distal from the bone (**Figure 6H**, **I**). Taken together, these findings suggest that AML cells at different states of differentiation locate in distinct niches within the BM.

**Figure 6:**
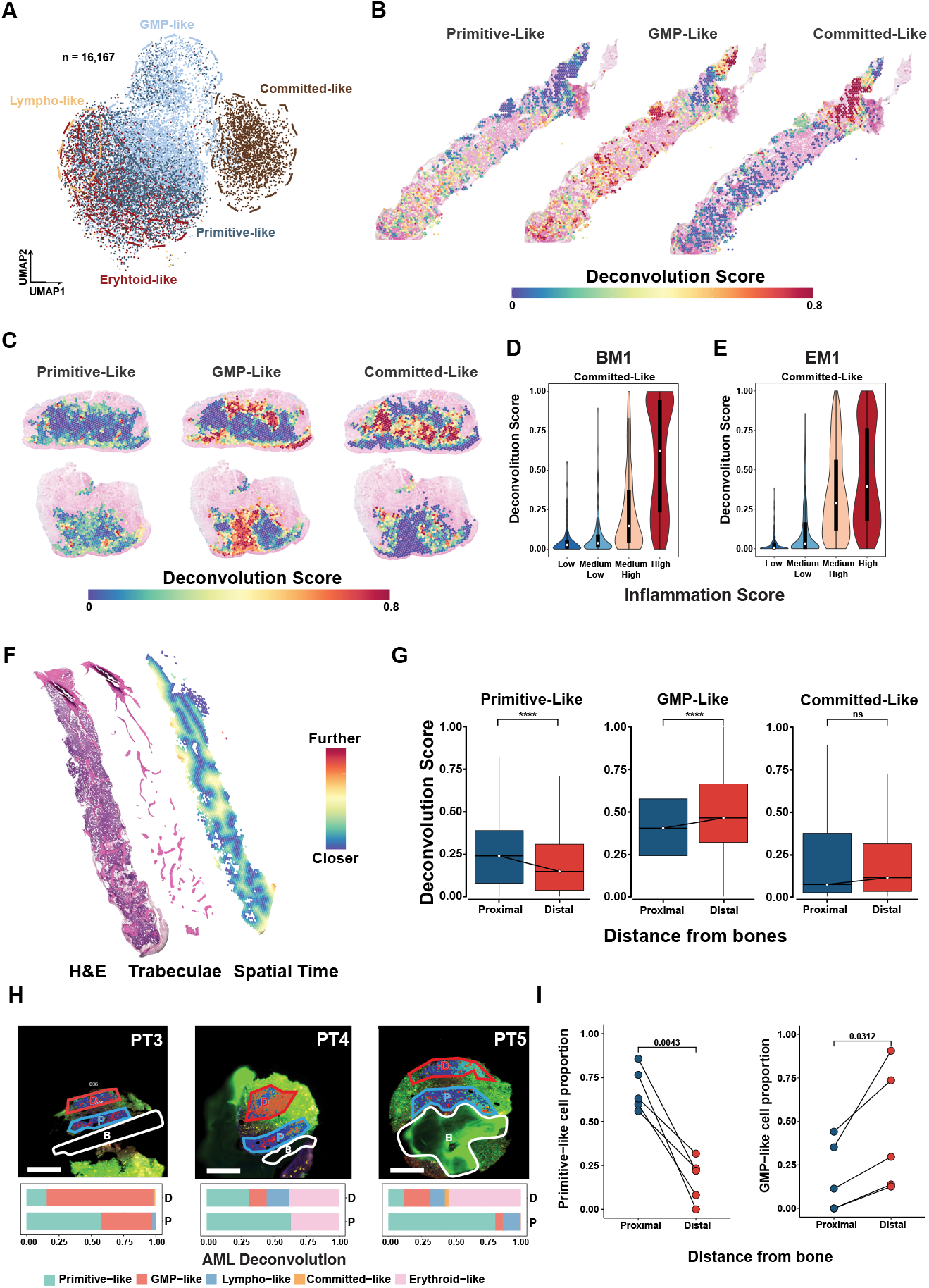
Localization of AML cell differentiation states within inflammatory and endosteal niches. **(A)** UMAP projection of 16,167 AML cells into differentiation states: primitive-like, granulocyte- monocyte progenitor (GMP)-like, erythroid-like and lymphoid-like, and committed-like (**B**) Spatial deconvolution maps of HLS-spots of BM1 showing primitive-like, GMP-like, and committed-like AML cells. **(C)** Spatial deconvolution maps of EM1 showing the same AML cell types. (**D, E)** Violin plot showing the distribution of committed-like AML cells in BM1 (**D**), and EM1 (**E**) across inflammation class. (**F)** Spatial map of Spatial Time calculation according to trabeculae overlaid with H&E image. (**G)** Box plots showing deconvolution scores of primitive-like, GMP-like, and committed-like AML cells relative to their distance from bone in Visium data. Proximal is dark blue, distal is dark red. **(H)** GeoMx analysis of AML deconvolution in bone marrow regions from 3 AML patients. D: distal (dark red), P: proximal (dark blue), B:bone (white). Stacked bar plots represent cell type deconvolution within distal and proximal regions. The scale bar is 250µm. **(I)** Line graphs showing proportions of primitive-like and GMP-like cells relative to distance from bone. *p < 0.05, **p < 0.01, ***p < 0.001, ****p < 0.0001, Wilcoxon rank sum test.

## Discussion

Understanding AML microenvironments requires unveiling the spatial niches along the medullary-extramedullary leukemia axis and the interactions between myeloid neoplastic populations and the immune system. Spatial transcriptomics (ST) technologies offer valuable insights but face challenges in BM-centric diseases due to preservation issues.^61,62^ In this study, we mapped human BM and EM tissues exhibiting the presence of AML using decalcified and archived core biopsy tissues, revealing significant interactions within the CXCL12-CXCR4 axis. This mapping highlights AML populations’ spatial localizations in various differentiation states within endosteal and inflammatory niches. Additionally, we validated these findings using GeoMX DSP and identified the most applicable Visium-based ST assays for leukemic tissues.

Inflammation is one of the hallmarks of cancer.^63^ Recent studies have shown that inflammatory states alter the immune microenvironment and are driven by specific differentiation stages in AML.^13,64–66^ These inflammatory microenvironments play roles in the maintenance of leukemic populations, disease progression, and chemoresistance.^67,68^ Here, inflammatory classification of spatial spots revealed niches in both AML-affected BM and EM leukemia. Although these inflammation niches harbored AML cells at various differentiation states, there was a strong association between a more committed-like phenotype and inflammation. The inflammation-high niches also created a hypoxic environment, linked to reactive oxygen species production and HIF pathway activation.^69^ Importantly, hypoxic microenvironments create protective niches that mediate resistance to therapy.^70^

Notably, we found that the inflammation niche was strongly associated with the region enriched for AML and monocytes, which communicate strongly through the CXCL12-CXCR4 axis. CXCL12, which is also a target of HIF-1α^71^, binds to CXC chemokine receptor 4 (CXCR4), a G-protein-coupled chemokine receptor, essential for the interaction between AML cells and the BM stroma. This interaction contributes to myeloid neoplastic progression, cell homing, and migration^72–75^ and has been studied as a therapeutic target in leukemia.^76–78^ CXCR4 mediates chemotaxis and retention of cells within the BM niche by interacting with the CXCL12 gradient.^79^ This axis is also thought to play a role in EM acute lymphoblastic leukemia and AML infiltration, with particularly higher expression levels identified in AML-M4/M5 subtypes.^80–82^ Here, we showed that spatial CXCR4 expression is highly abundant and widespread in EM tissue. Therefore, consistent with previous studies, it is possible that CXCR4 is not only a homing factor for the BM but also facilitates the migration and homing of leukemia populations to EM sites.

CXCR4 has previously been implicated in EMT-like transcriptional activation in EM disease in multiple myeloma.^83^ The PI3K/Akt/mTOR pathway is one of the downstream targets of the CXCL12-CXCR4 axis and is recognized as an important regulator of EMT in solid tumors, playing a crucial role in tumor cell migration and metastasis.^57,58,84,85^ In AML, this EMT-like pathway, which we refer to as a trans-differentiation state, shows correlation with the PI3K/Akt/mTOR pathway in inflammatory niches with high CXCL12-CXCR4 signaling. Targeting the CXCL12-CXCR4-enriched inflammatory niche may therefore reduce trans-differentiation and the development of leukemia cutis, offering potential benefits for patients with concurrent medullary and extramedullary AML.

The AML population in the BM exhibited a transcriptional profile similar to that of EM leukemia within this inflammatory niche, revealing both common and specific genes and pathways for spatial niche analysis. Beyond the previously identified chimeric antigen receptor (CAR)-T therapy target, *CD70*,^86,87^ we observed the upregulation of genes such as *RAB3D*,^88^ associated with monocytic AML subtypes, and *TP53INP2*,^52,89^ linked to autophagy activity in NPM1-mutated AML, within these regions. Additionally, the high expression of *TNFSF13B* and *TMEM176B*, markers not yet definitively linked to AML, in regions populated by monocyte-like AML populations and in corresponding EM leukemia areas, highlights the need for further research on these markers for prognosis and as potential therapeutic targets.

The interaction between hematopoiesis and the endosteal niche in the BM is crucial for maintaining HSC quiescence, self-renewal, and differentiation; facilitating HSC homing and mobilization, and supporting the BM microenvironment.^90,91^ Previous studies have shown that HSCs and multipotent progenitors are closely associated with the bone surface in the endosteal region, whereas committed progenitors and differentiated cells are positioned more distally.^92–95^ Additionally, a comprehensive spatial atlas of the human BM niche has been provided by cellular biogeography mapping studies.^96^ Here, we extended prior findings by examining spatial interactions in BM with AML. The differentiation states of AML cells are challenging to define using individual markers. We provided a new approach to examine the connection between AML populations at distinct differentiation states and the endosteal niche in BM tissues using multimodal spatial transcriptomic analysis. Our findings suggest that while leukemic populations in the GMP and committed differentiation states are located further from the endosteal niche, primitive-like cells are in closer proximity. This suggests that leukemic cell maintenance and stemness are supported by this niche.

In summary, our study provides a comprehensive ST analysis of medullary and EM tissues in AML using multimodal ST technologies. We identified distinct inflammatory niches in both BM and EM samples, with significant involvement of the CXCL12-CXCR4 axis. This axis promotes trans-differentiation via the PI3K/AKT/mTOR pathway, contributing to AML cell migration and infiltration into EM sites. Spatial analysis revealed that primitive-like AML cells are predominantly located near the endosteal niche in BM, while committed-like and GMP-like populations are situated more distally. These findings highlight the critical role of inflammatory niches in AML progression and provide novel insights into leukemic cell spatial heterogeneity, underscoring potential targets for therapeutic intervention.

## Limitations of the study

Our study presents one of the initial spatial analyses at the RNA level along the human medullary-extramedullary leukemia axis, yet is limited by the number of samples with only male cohort. This is due in part to the cost-prohibitive nature of these assays (Visium, mfIHC and DSP). Nevertheless, our study represents the first study evaluating the spatial dynamics of medullary and EM leukemia in paired patient samples constituting a very rare sample population, which will serve as a resource for other researchers. mfIHC was performed on tissue sections that were not immediately adjacent to those used for ST which may introduce minor discrepancies in spatial alignment. Our in-house generated scRNA-seq reference dataset was prepared from bone marrow mononuclear cells (BMMCs), which excludes neutrophils due to their higher density and susceptibility to degradation during isolation procedures. Additionally, the 55-µm spot-based structure of Visium ST technology leads to low resolution in tissues like BM, which have highly heterogeneous cellular compositions. To overcome this limitation, we have implemented several strategies: (i) We adapted a MAD-based filtering method specific to each tissue structure to identify spots with outlier reads. (ii) We defined intra-sample niches using broader range clusters. (iii) We validated cell prediction results with pathology annotations made using both H&E and immunohistochemical analysis. (iv) We validated the Visium results using DSP and mfIHC for higher resolution.

## Code Availability

Code for figure generation and key analysis are available at http://github.com/abbaslab/

## Data Availability

Visium spatial transcriptomics data will be publicly available at GEO with accession number GSE279576 as of the date of publication. Additionally, an interactive web application will be launched concurrently, allowing researchers to explore the data interactively.

## Acknowledgements

We would like to thank Sarah Bronson at the Research Medical Library and Thomas Huynh and Arizona Nguyen at the Department of Veterinary Services for assistance in histologic sample preparation, and Dr. Chong Wu at the Department of Biostatistics at The University of Texas MD Anderson Cancer Center for reading and providing comments on the manuscript. This work was supported in part by the University of Texas MD Anderson Cancer Center Support Grant CA016672 and the University of Texas MD Anderson Cancer Center Physician-Scientist Training Program, philanthropic funding and MG from Energy Transfer, and a donation to HAA from the Diego-Osio Llerenas Fund. HAA was funded by the Physician Scientist Award and Cancer Prevention and Research Institute of Texas (CPRIT).

## Contributions

H.A.A. conceived the study, supervised all aspects of the work, co-wrote and reviewed the manuscript. E.D. led the computational analyses and interpretation of Visium data. E.D., I.V. and H.A.A. wrote the manuscript. I.V. and C.P.L. analyzed DSP data. C.D.P. applied and analyzed the mfIHC. A.E.Q. and F.Z.J. performed pathology annotations. P.B., S.B., S.J., Z.W., A.L., and K.M.W. conducted the experiments and/or library preparation. D.A.A., P.H.G., P.K.R., P.S., and R.J.T., contributed conceptually to data analysis and design. All authors read and edited the manuscript.

---

**Figure S1: Evaluation of RNA and DNA Integrity, and Spatial Transcriptomics Quality Metrics for Bone Marrow and Extramedullary Tissues from AML Patients**

**(A)** Histological sections of the BM1, BM2, EM1, and EM2 samples with hematoxylin and eosin (H&E) staining. Each section shows distinct tissue architecture and regions. Annotation based on pathology annotation. Scale bars represent 1 mm. (**B**) H&E-stained section showing necrotic phenotype in the EM2 sample. While black scale bar at main tissue reprent 1 mm, zoomed-in white scale bar represent 50 µm. **(C)** Sections of the EM1 sample. Scale bars indicate 1 mm. (**D)** Pre-library RNA fragmentation traces (DV_200_ values) for the four samples. EM2 showed the lowest DV_200_ at 24%, while BM2 had the highest at 62% **(E)** Post-library DNA traces for EM tissues comparing Visium v1 and v2 assays. Similar DNA trace profiles were observed between v1 and v2 assays for both EM samples. (**F)** Post-library DNA traces for bone marrow tissues comparing Visium v1 and v2 assays. V2 assays displayed significantly higher-post-library DNA traces compared to off-peaks at v1 assay. (**G**) Comparison of mean reads under tissue per spot between Visium v1 and v2 assays for all four samples. **(H)** Median genes per spot detected in Visium v1 and v2 assays for all four samples. **(I)** Percentage of reads mapped confidently to the filtered probe set for Visium v1 and v2 assays across the samples. **(J)** Correlation between median number of oligonucleotides and median gene detected, showing a strong positive correlation (r = 0.996, p = 1.4 × 10^-5^). **(K)** Spatial distribution and comparison of mitochondrial gene reads (%) at the spots at the edge versus others. **(L)** Radar plot comparing DV200 values and median mitochondrial gene reads (%) for the four sample (**M**) Histological overlay of an H&E-stained BM2 section with the CytAssist image and ST data reveals preserved structural integrity of bone regions with black lines. Scale bar indicate 1 mm.

**Figure S2: Median Absolute Deviation (MAD) Filtering and Spot-Level Quality Metrics in ST analysis**

**(A)** Workflow of Visium ST processing for v1 and v2 assays, including alignment, transcriptome counting, and subsequent quality control steps. (**B**) Mitochondrial content distribution across the four samples (BM1, BM2, EM1, EM2). Scatter plots show mitochondrial gene read percentages against the number of genes detected per spot, with MAD filtering applied (red dashed lines). **(C)** Expression profile distributions for the four samples. Scatter plots depict log-transformed counts of genes and oligonucleotides per spot, with MAD filtering indicated by red dashed lines. **(D)** Spatial distribution of mitochondrial gene content and MAD-filtering results for BM1, BM2, EM1, and EM2 samples. Insets highlight regions with filtered spots. **(E)** Bar graph showing the percentage of spots removed after applying MAD filtering based on mitochondrial content and expression profile for each sample. (**F)** Comparison of removed spots between different sections of the EM1 sample based on mitochondrial content and expression profile.

**Figure S3: Pathology Annotation and validation of deconvolution**

**(A)** Hematoxylin and eosin (H&E)-stained images and CytAssist images for BM1, BM2, EM1, and EM2 samples. Scale bars represent 1 mm. (**B**) Ground truth, spatial clusters, and UMAP representations of the BM1, BM2, EM1, and EM2 samples. Unsupervised clustering segmented the tissues into distinct clusters. **(C)** Shared nearest neighbor (SNN) graph-based clustering of BM1, showing regions and unsupervised clusters. (**D)** SNN graph-based clustering of BM2, showing pathology clusters and unsupervised clusters. **(E)** Detailed pathology annotation for BM1, showing regions annotated as erythroid-enriched, leukemic tissue, and other areas. Sankey diagram illustrates the overlap between detailed pathology annotations and identified regions. (**F)** Immunohistochemical staining for CD11c, MPO, and CD3e on BM1 sections. The scale bar for the main tissue panels represents 1 mm. The scale bar for the zoomed-in panels, corresponding to the boxed regions, represents 100 µm. (**G)** Dot plot showing the cell type deconvolution in bone marrow samples, highlighting differences in cell population in each sample. **(H)** Dot plot showing the cell type deconvolution in extramedullary samples, highlighting differences in cell population in each sample. **(I)** Heatmap showing the differential expressed genes of EM1, annotated by unsupervised clusters.

**Figure S4: Inferred Pathway Analysis and Relationship with Inflammation in AML Bone Marrow and Extramedullary Tissues**

**(A)** Differential deconvolution analysis between high leukemic score (HLS) and low leukemic score (LLS) spots in BM1. (**B**) AUCell score distribution of HLS spot genes across 3 clusters, highlighting the concentration of HLS genes in cluster 3. **(C)** Heatmap of pathway activities in BM1, BM2, EM1, and EM2 samples. Hierarchical clustering show same patients’ sample similarities. (**D)** Spatial clustering of pathway activities in BM1, EM1, BM2, and EM2 samples. Unsupervised clusters and corresponding pathway profiles are shown, indicating the similarity between cluster 3 of BM1 and leukemic population cluster of EM1. **(E)** Inflammation score distribution across spatial clusters in BM1. High inflammation profile is concentrated in cluster 3. **(F)** Inflammation score distribution across spatial clusters in EM1, highlighting high inflammation at leukemic infiltration clusters. (**G)** Spatial dynamics of inflammation score calculated with SpatialTime, indicating elevated Hypoxia, Reactive Oxygen Species, and HIF1A pathways near high inflammatory spots. **(G)** Dot plot for T cell subset based on inflammation class in PT2 samples.

**Figure S5: Spatial Cell-Cell Interactions and Pathway Analysis of CXCL12-CXCR4 Axis in AML Bone Marrow and Extramedullary Tissues**

**(A)** Spatial map of BM1 with spots labeled by cell type based on deconvolution results, highlighting the distribution of different cell populations. (**B**) Heatmap of outgoing and incoming signaling patterns for various cell types in BM1, showing strong signaling activity in CXCL pathways. **(C)** Chord diagram illustrating the interactions between CXCL12 and CXCR4 across different cell types in BM1. **(D)** Dot plot of communication probabilities for significant ligand-receptor pairs, between AML-GMP-Monocytes showing strong interactions involving CXCL12- CXCR4 in BM1. **(E)** Correlation of CXCL signaling with other hallmark pathways, highlighting significant associations with inflammation-related pathways. Specifically, mTORC1 and PI3K/AKT/mTOR pathways highly correlated. (**F)** Violin plot of PI3K/AKT/mTOR signaling activity in BM1, stratified by inflammation score categories, indicating higher pathway activity in regions with elevated inflammation. (**G)** Scatter plot showing the correlation between the composite inflammation score and epithelial-mesenchymal transition (EMT) markers in BM1. **(H)** Violin plot of PI3K/AKT/mTOR signaling activity in EM1, stratified by inflammation score categories. **(I)** Violin plot of epithelial-mesenchymal transition (EMT) markers in EM1, stratified by inflammation score categories.

**Figure S6: Multiplexed fluorescence immunohistochemistry aligns with ST signatures**

**(A)** mfIHC on near-adjacent tissue sections from Visium samples EM1, BM2, and EM2, showing merged phenotypic (CD33, CD68, and CD71) and functional (CXCL12, CXCR4, and IL-6) markers. (**B**) Spatial distribution of phenotypic and **(C)** functional markers on EM1 sample. Zoomed OPAL and Visium images demonstrating concordance. **(D, E)** Phenotypic and functional marker staining for BM2 and EM2 shows similar patterns with ST data. (**F)** mfIHC IL-6 intensity level shows correlation with composite score comes from ST data. (Pearson correlation) (**G)** Comparison of CXCL12 intensities in inflammatory regions across BM1 and EM1, with enrichment in high inflammation zones in BM1. Scale bars: 1 mm (whole-slide panels) and 100 µm (selected region panels).

**Figure S7: Classification and Spatial Distribution of AML Cells Based on Differentiation States**

**(A)** Spatial map of BM1 showing high leukemic spots (HLS) divided by unsupervised clusters. (**B**) Spatial distribution of different AML cell state deconvolution in BM1, classified as primitive-like, GMP-like, committed-like, erythroid-like, and lymphoid-like. **(C)** Density plots showing the distribution of primitive-like and committed-like populations across unsupervised clusters in BM1. **(D)** Bar graph of deconvolution scores for primitive-like and committed-like cells in BM1, indicating a higher abundance of committed-like cells. **(E)** Bar graph of deconvolution scores for primitive-like and committed-like cells in EM1, showing a reduced primitive-like population in EM tissue. (**F)** Spatial map of BM1 indicating the localization of spots relative to trabecular bone regions. Proximal: Dark blue and Distal: Dark red (**G)** Immunofluorescence image of the GeoMx RNA assay with ROI labels. AOIs captured by GeoMx segmented by marker expression are shown within each ROI (CD34 enriched in green, CD68 enriched in red, DNA in blue). **(H)** Immunofluorescence image of the GeoMx protein assay with ROI labels. AOIs captured by GeoMx segmented by marker expression are shown within each ROI (CD34 enriched in green, CD68 enriched in red, DNA in blue). **(I)** Raw RNA counts of the DNA GeoMx assay. Red lines demark the limit of quantification (LOQ) calculated for each AOI. Pie charts at the bottom show the percentage of genes above the LOQ per segment. **(J)** Deconvolution of GeoMx RNA AOIs using signatures derived from scRNA dataset. Primitive-like: HSC-like, CMP/LMPP-like; Lymphoid-like: CLP-like, Lymphoid-like; Committed-like: Mono-like, Basophil-like, DC-like.

