## Supplementary figures and images for "Spatial Transcriptomics Reveals Inflammation and Trans-differentiation States of Acute Myeloid Leukemia in Extramedullary and Medullary Tissues"

### Supplementary Figure 1

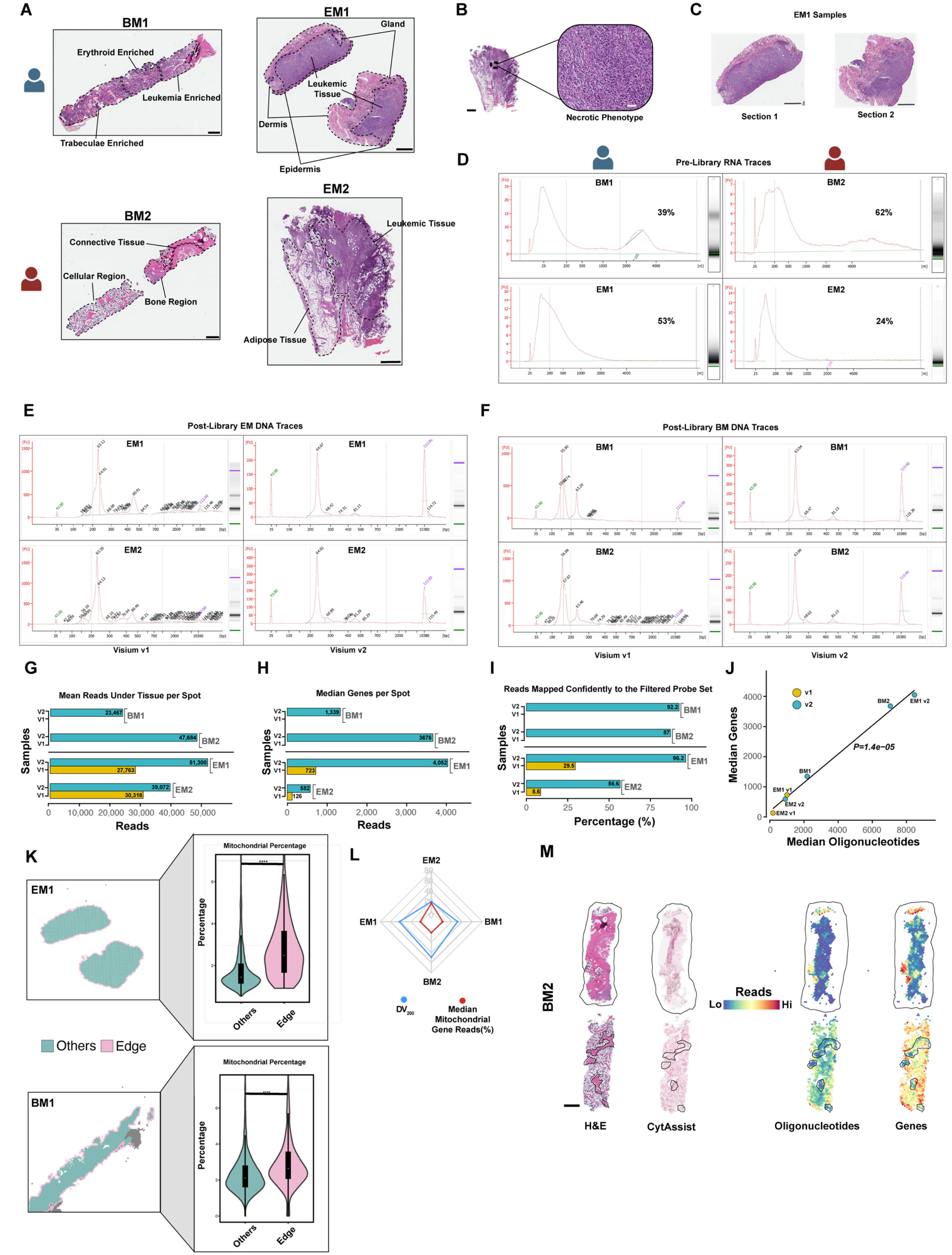

### Supplementary Figure 2

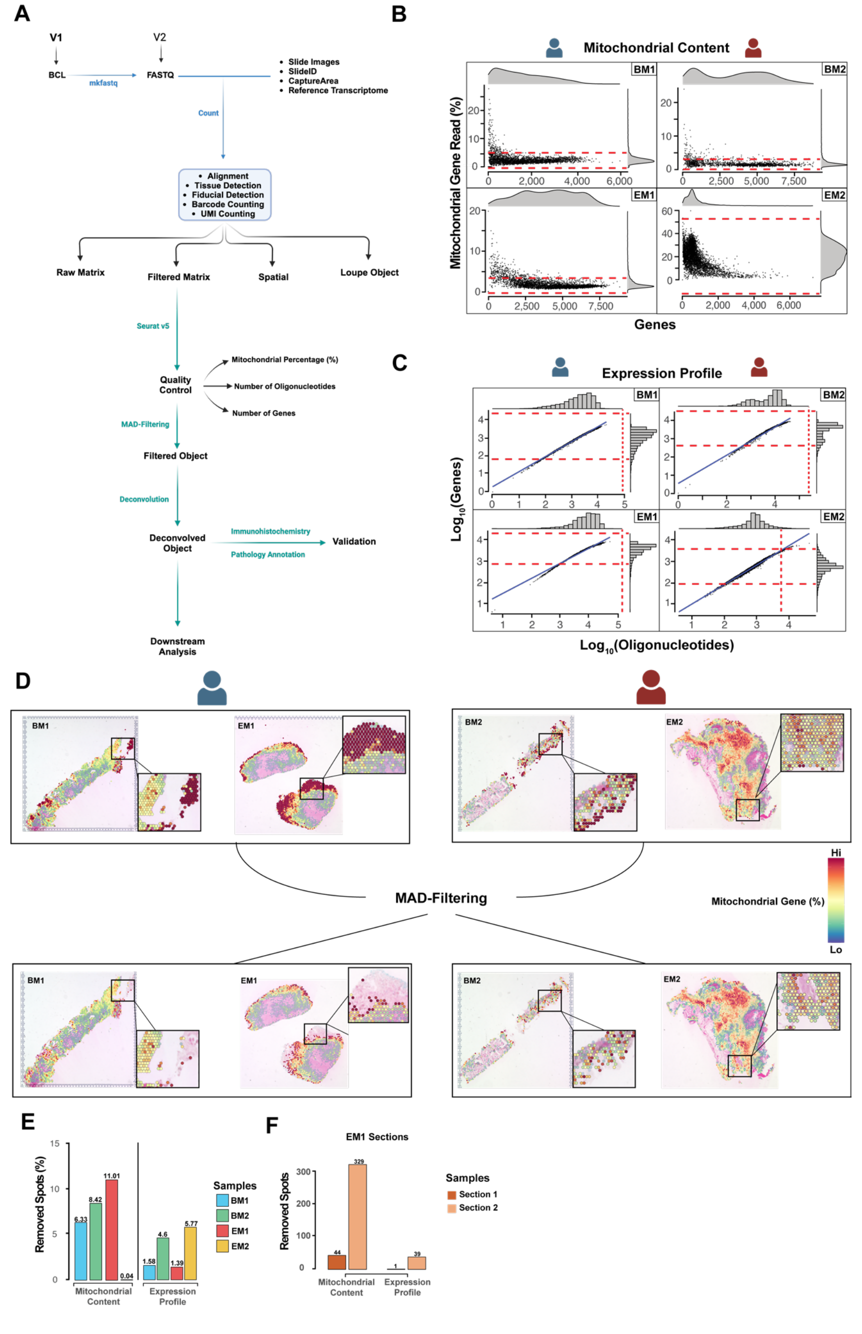

### Supplementary Figure 3

#### Supplementary Figure 3

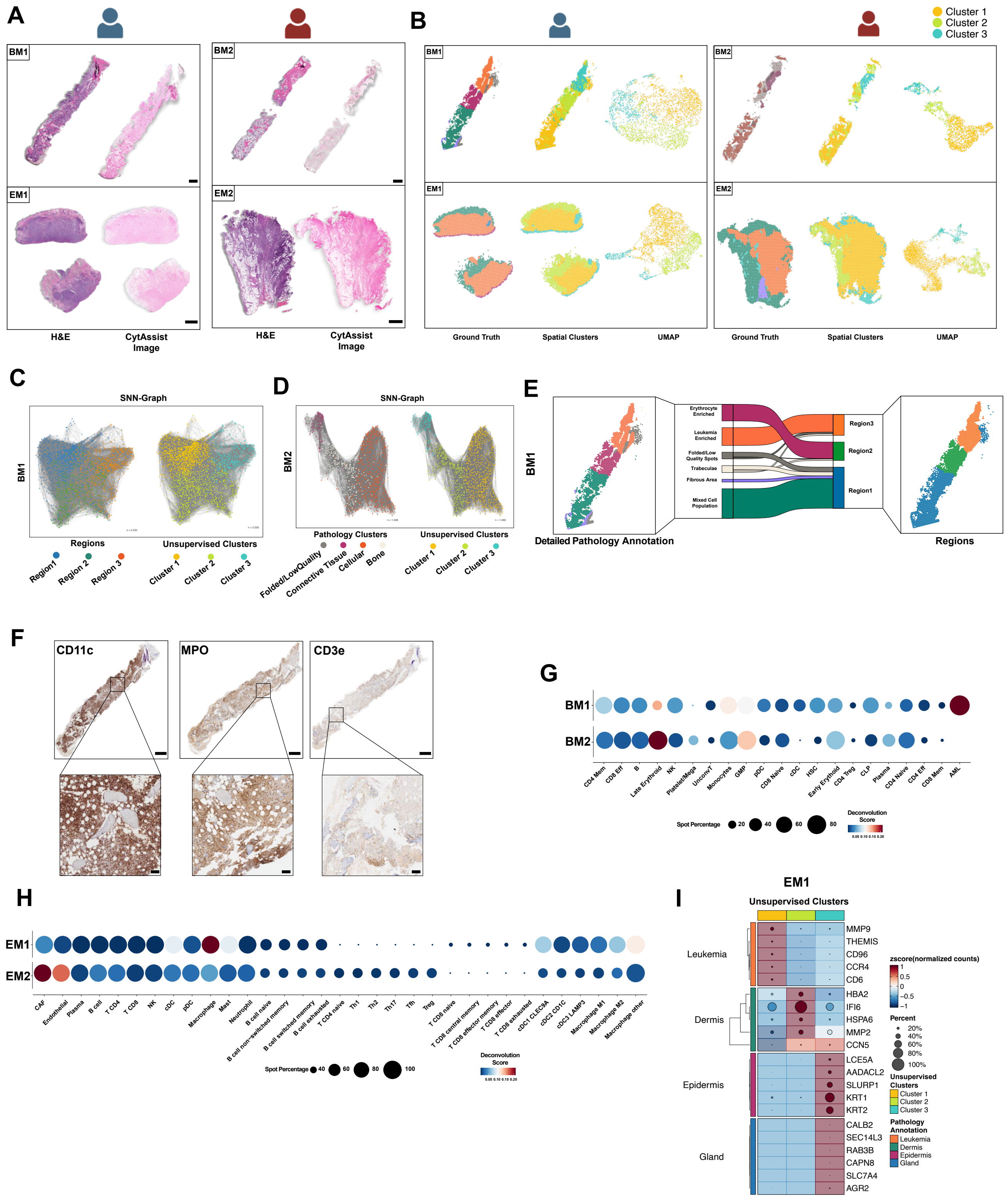

### Supplementary Figure 4

#### Supplementary Figure 4

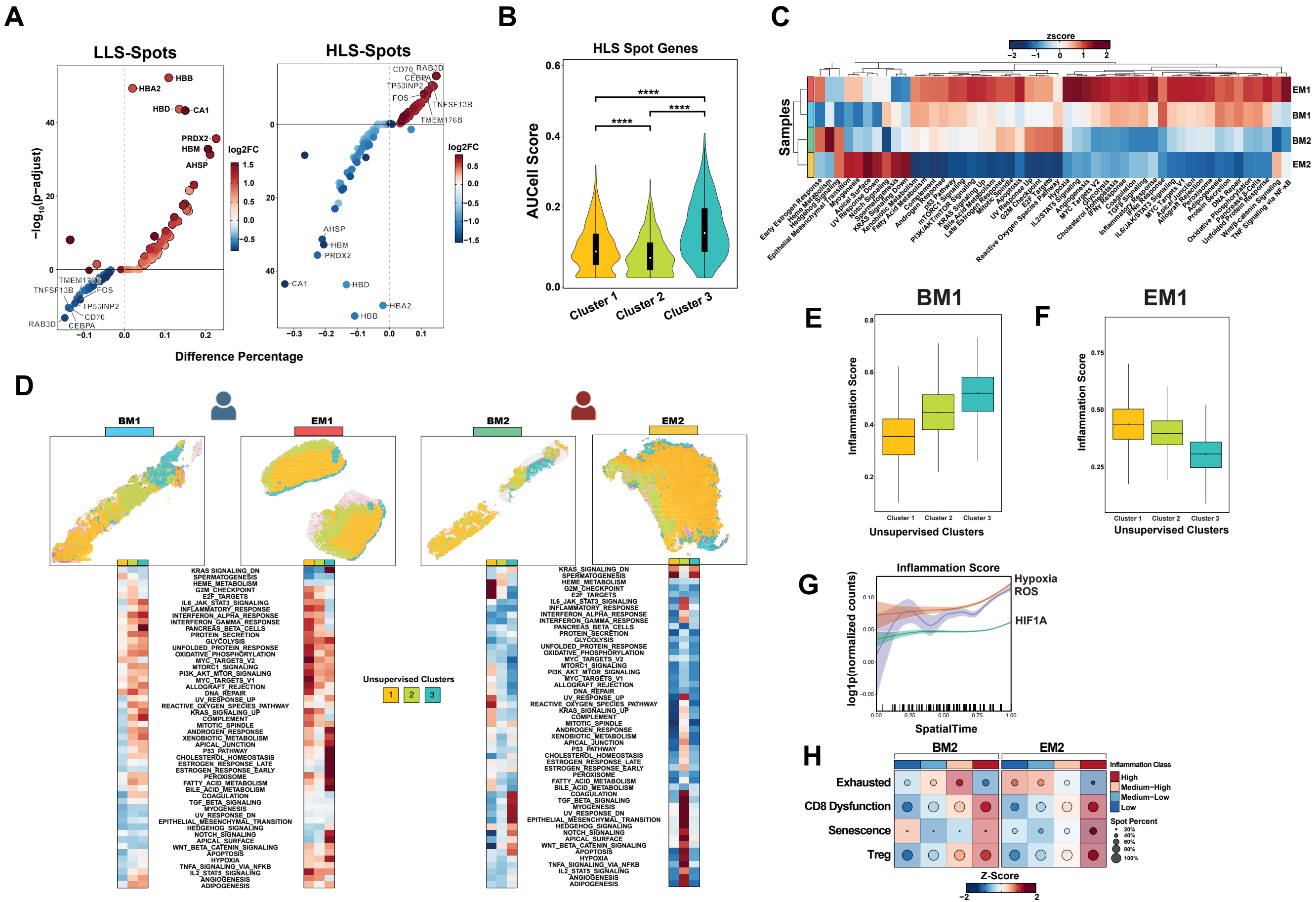

### Supplementary Figure 5

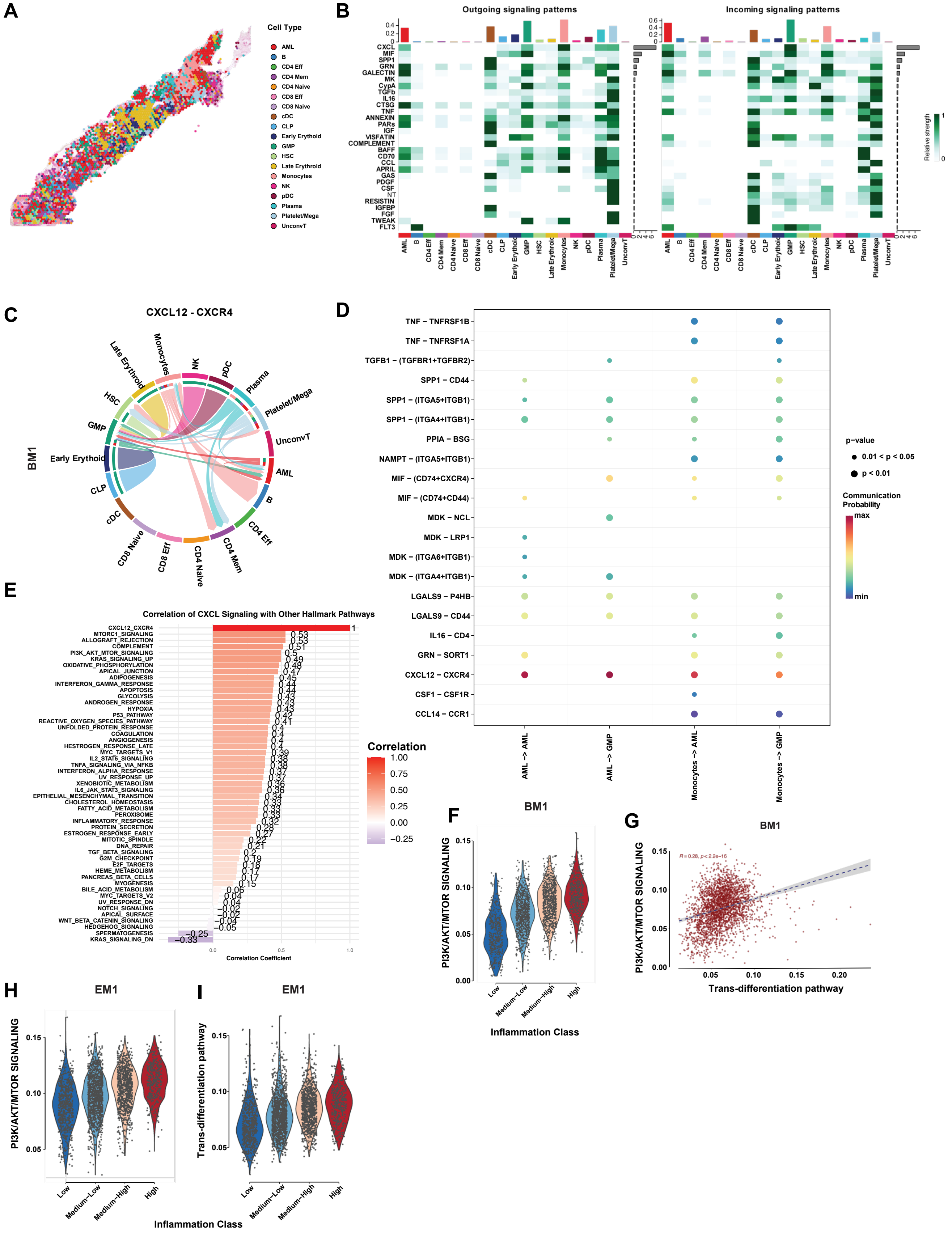

### Supplementary Figure 6

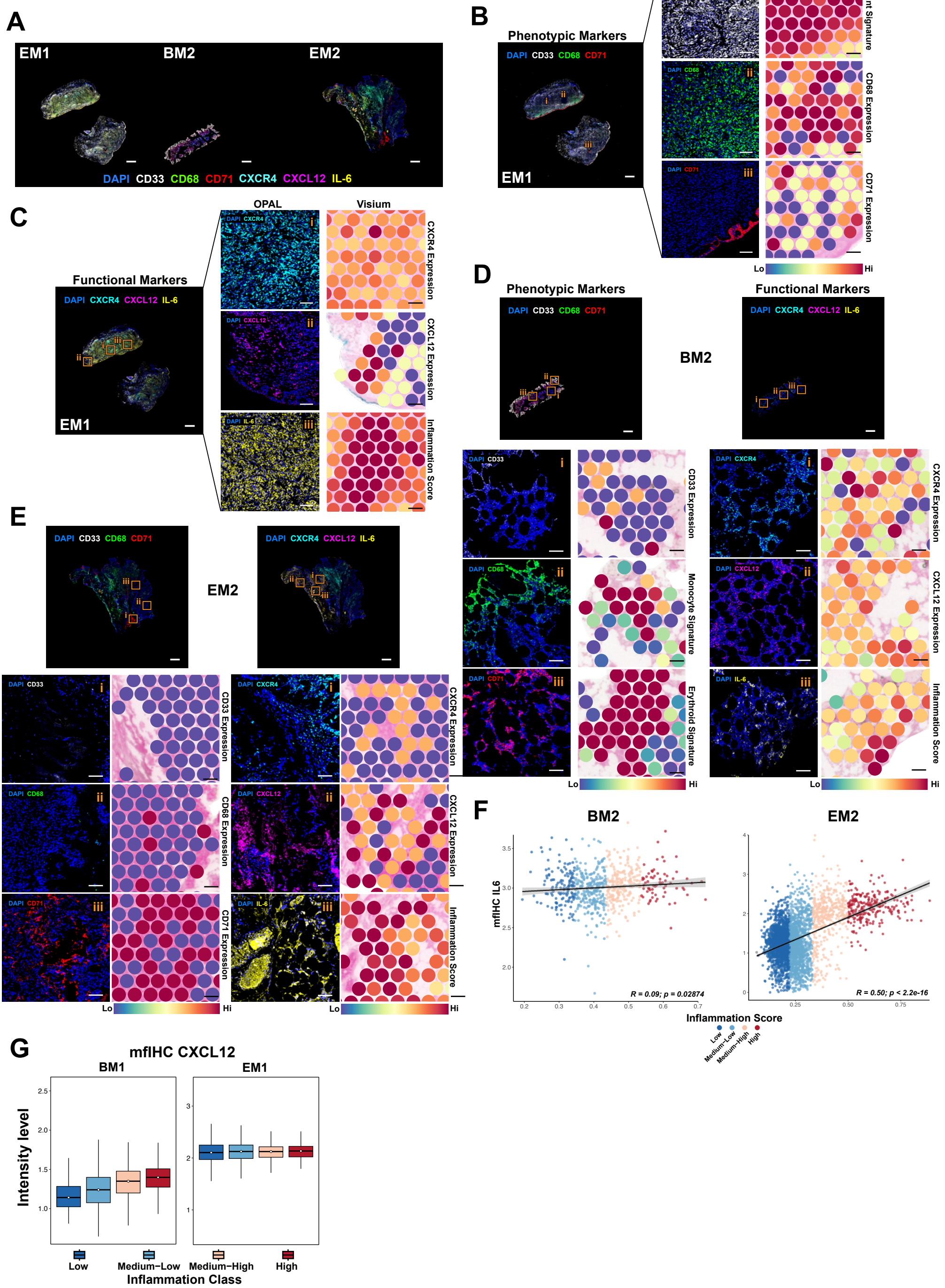

### Supplementary Figure 7

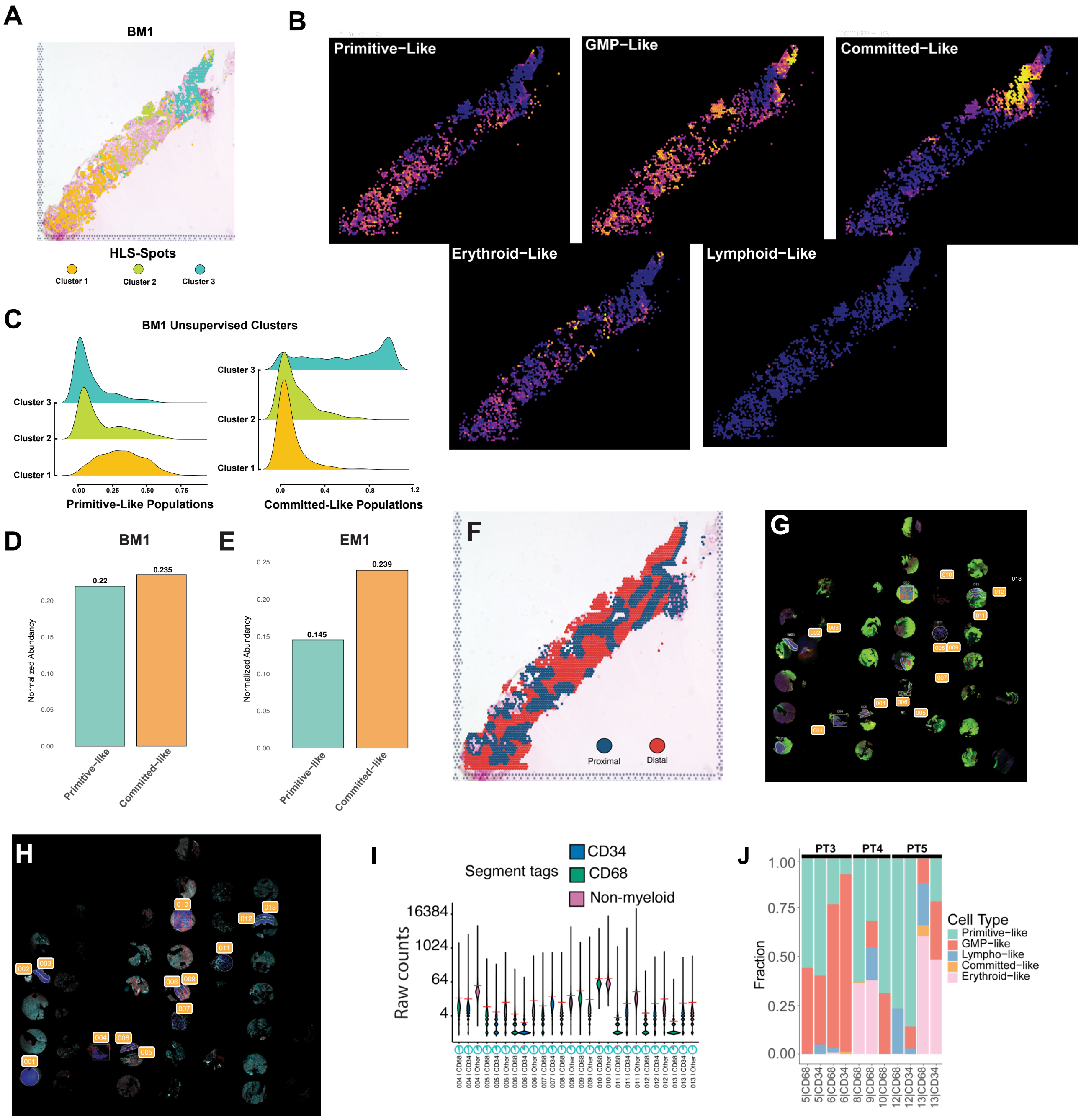
