## Supplemental Table 1 for "Spatial Transcriptomics Reveals Inflammation and Trans-differentiation States of Acute Myeloid Leukemia in Extramedullary and Medullary Tissues"

**Table S1. Patients’ characteristics**

| **Characteristic (unit)** | **PT1** | | | **PT2** | **PT3** | | | **PT4** | | **PT5** |
| --- | --- | --- | --- | --- | --- | --- | --- | --- | --- | --- |
| **Demographics** | | | | | | | | | | |
| Age (years) | 42 | | | 83 | 70 | | | 75 | | 74 |
| Sex/gender | Male | | | Male | Male | | | Male | | Male |
| Race/ethnicity | Hispanic | | | Hispanic | White | | | White | | Asian |
| **Disease progression** | | | | | | | | | | |
| AML subtype^1^ | EM-AML/MS | | | EM-AML/MS | AMoL/AML-M5 | | | AML-M1 | | AML-MRC |
| Origin | De novo | | | De novo | De novo | | | De novo | | sMDS |
| Time since diagnosis (months) | 0 | | | 0 | 0 | | | 0 | | 0 |
| Risk category^2^ | Favorable | | | Adverse | Adverse | | | Adverse | | Adverse |
| Status | Deceased | | | Deceased | Deceased | | | Alive | | Deceased |
| Survival time (months) | 2 | | | 1 | 4 | | | 67 | | 8 |
| **Spatial Transcriptomic sample information** | | | | | | | | | | |
| Disease status | | Newly Diagnosed | Newly Diagnosed | | | Newly Diagnosed | Newly Diagnosed | | Newly Diagnosed | |
| Collected year | | 2018 | 2017 | | | 2015 | 2018 | | 2017 | |
| **Data** | | | | | | | | | | |
| Technology | | Visium | Visium | | | GeoMx | GeoMx | | GeoMx | |
| Assay | | v1 + v2 | v1 + v2 | | | Hu WTA | Hu WTA | | Hu WTA | |
| **Cytogenetic and molecular features** | | | | | | | | | | |
| Karyotype | Diploid | | | Diploid | Monosomy 7 | | | Diploid | | Monosomy 7 |
| Mutations^3^ | None detected | | | DNMT3A, FLT3, IDH1, IDH2, KRAS, NPM1, RAS, SF3B1 | DNMT3A, IDH1 | | | IDH2, NPM1, SRSF2 | | ASXL1, EZH2, KIT, NRAS, RUNX1 |
| **Bone marrow counts** | | | | | | | | | | |
| Cellularity (%) | 40-50 | | | 80-90 | 90-100 | | | 90-100 | | 90-100 |
| Myeloblasts (%) | 2 | | | 30 | 52 | | | 65 | | 31 |
| Monocytes (%) | 2 | | | 3 | 17 | | | 1 | | 2 |
| **Peripheral blood counts** | | | | | | | | | | |
| Leukocytes (×10^9^/L) | 6.2 | | | 11.1 | 8.3 | | | 17.1 | | 25.1 |
| Myeloblasts (%) | 0 | | | 8 | 13 | | | 80 | | 58 |
| Monocytes (%) | 8 | | | 6 | 51 | | | 1 | | 7 |
| **Treatment** | | | | | | | | | | |
| Active agents | CLIA | | | DAC | DAC | | | CL+LDAC+VEN  / AZA+VEN | | CL+LDAC |
| Response category | NR | | | NR | NR | | | CR | | NR |

Abbreviations: EM, extramedullary; MS, myeloid sarcoma; AMoL, acute monocytic leukemia; MRC, myelodysplasia-related changes; sMDS, secondary to myelodysplastic syndrome; v1, Version 1; v2, Version 2; Hu WTA, Human Whole Transcriptome Atlas; CLIA, cladribine, idarubicin, and cytarabine; DAC, decitabine; CL, cladiribine; LDAC, low-dose cytarabine; VEN, venetoclax; AZA, azacytidine; NR, no response; CR, complete remission.

^1^ AML subtypes were determined according to the World Health Organization’s 5th edition [96].
^2^ Risk categories were estimated using the European LeukemiaNet 2022 guidelines [97].
^3^ Mutations were detected using the 81-gene next-generation sequencing panel, EndLeukemia Assay v1 [98].
